# Mosaic and non-mosaic protocadherin 19 mutation leads to neuronal hyperexcitability in zebrafish

**DOI:** 10.1101/2021.09.03.458732

**Authors:** Barbara K. Robens, Xinzhu Yang, Christopher M. McGraw, Laura H. Turner, Carsten Robens, Summer Thyme, Alexander Rotenberg, Annapurna Poduri

**Affiliations:** Department of Neurology, F.M. Kirby Neurobiology Center, Boston Children’s Hospital – Harvard Medical School, Boston, MA, USA; Epilepsy Genetics Program, Department of Neurology, Boston Children’s Hospital – Harvard Medical School, Boston, MA, USA; Department of Neurology, Harvard Medical School, Boston, MA, USA; Division of Epilepsy, Department of Neurology, Massachusetts General Hospital, Boston, MA, USA; MIT-Harvard Center for Ultracold Atoms, Research Laboratory of Electronics, and Department of Physics, Massachusetts Institute of Technology, Cambridge, Massachusetts 02139, USA; Department of Neurobiology, University of Alabama at Birmingham, Birmingham, AL, USA; Division of Epilepsy and Clinical Neurophysiology, Department of Neurology, Boston Children’s Hospital, Boston, MA, USA

**Keywords:** PCDH19, girls clustering epilepsy, hyperexcitability, zebrafish, mosaicism, epilepsy, X-linked, anxiety, learning and memory, inhibitory neurons

## Abstract

Epilepsy is one of the most common neurological disorders. The X-linked gene *PCDH19* is associated with sporadic and familial epilepsy in humans, typically with early-onset clustering seizures and intellectual disability in females but not in so-called ‘carrier’ males, suggesting that mosaic PCDH19 expression is required to produce epilepsy. To characterize the role of loss of PCDH19 function in epilepsy, we generated zebrafish with truncating *pcdh19* variants. Evaluating zebrafish larvae for electrophysiological abnormalities, we observed hyperexcitability phenotypes in both mosaic and non-mosaic *pcdh19*^+/-^ and *pcdh19*^-/-^ mutant larvae. Thus, we demonstrate that the key feature of epilepsy—network hyperexcitability—can be modeled effectively in zebrafish, even though overt spontaneous seizure-like swim patterns were not observed. Further, zebrafish with non-mosaic *pcdh19* mutation displayed reduced numbers of inhibitory interneurons suggesting a potential cellular basis for the observed hyperexcitability. Our findings in both mosaic and non-mosaic *pcdh19* mutant zebrafish challenge the prevailing theory that mosaicism governs all PCDH19-related phenotypes and point to interneuron-mediated mechanisms underlying these phenotypes.

## Introduction

Epilepsy is a common chronic disorder of the brain characterized by recurrent, unprovoked seizures with a substantial contribution from genetic causes. The X-linked gene *PCDH19*, encoding the cell-cell adhesion molecule protocadherin 19, has emerged as one of the most prominent single genes associated with epilepsy (Dibbens et al., 2008; Duszyc et al., 2015; Epi4K Consortium and Epilepsy Phenome/Genome Project, 2017). Pathogenic variants in *PCDH19* were shown to cause PCDH19-clustering epilepsy (CE), previously referred to as Epilepsy and Mental Retardation Limited to Females (EFMR), OMIM #300088). Disease manifestations vary in severity, even among individuals with the same variant. They include early-onset seizures that are often fever-related and cluster with multiple seizures occurring one after another, intellectual disability, and autism (Dibbens et al., 2008; Dimova et al., 2012; Smith et al., 2018). The disorder follows a unique inheritance pattern, affecting primarily heterozygous females and sparing hemizygous males, at least from overt epilepsy and severe symptomatology; rarely observed males who are mosaic for hemizygous variants in this gene also display these major symptoms (Kolc et al., 2018). It has been proposed that the tissue mosaicism caused by random X-inactivation in all female cells leads to an abnormal cellular pattern during development. The so-called ‘cellular interference’ hypothesis posits that the presence of a mixture of cells in the brain, expressing either wild-type or mutant PCDH19 protein, leads to abnormal brain development and is consequently the underlying cause of the clinical manifestations (Depienne et al., 2009). However, it is still unclear how mosaic expression of PCDH19 in the brain leads to epileptogenesis and whether mosaic expression of wild-type (WT) and mutant protein in neurons is the sole cause for the clinical manifestations of PCDH19-CE.

The *PCDH19* gene consists of 6 exons, and most of the over 100 identified patient variants occur in exon 1, which encodes the 6 extracellular cadherin domains (Figure 1) (Kolc et al., 2018). PCDH19 is highly expressed in the brain and belongs to the largest subgroup of cadherins that are involved in calcium-dependent cell-cell adhesion (Depienne and LeGuern, 2012). However, the exact function of PCDH19 is still largely unknown. Recent studies in mice suggest that PCDH19 determines cell adhesion affinities and is involved in cell sorting during cortical development and mossy fiber synapse development (Hoshina et al., 2021; Pederick et al., 2018). Even though *Pcdh19* heterozygous mice do not show gross behavioral abnormalities or spontaneous seizures, treatment with the γ-aminobutyric acid type A (GABA-A) receptor inhibitor flurothyl revealed increased seizure susceptibility; interestingly, this finding does not require tissue mosaicism as it is present in both female heterozygous and homozygous mice (Hayashi et al., 2017; Pederick et al., 2016; Pederick et al., 2018; Rakotomamonjy et al., 2020). Other *in vitro* studies suggest a role for Pcdh19 in establishing proper brain architecture and neuronal connections (Mincheva-Tasheva et al., 2021; Pederick et al., 2016). Similar roles have been proposed for the highly conserved zebrafish (*Danio rerio*) Pcdh19 protein which was shown to be involved in cell proliferation and neuronal organization of the optic tectum, the largest region of the zebrafish brain, important for visual processing (Cooper et al., 2015).

**Figure 1:**
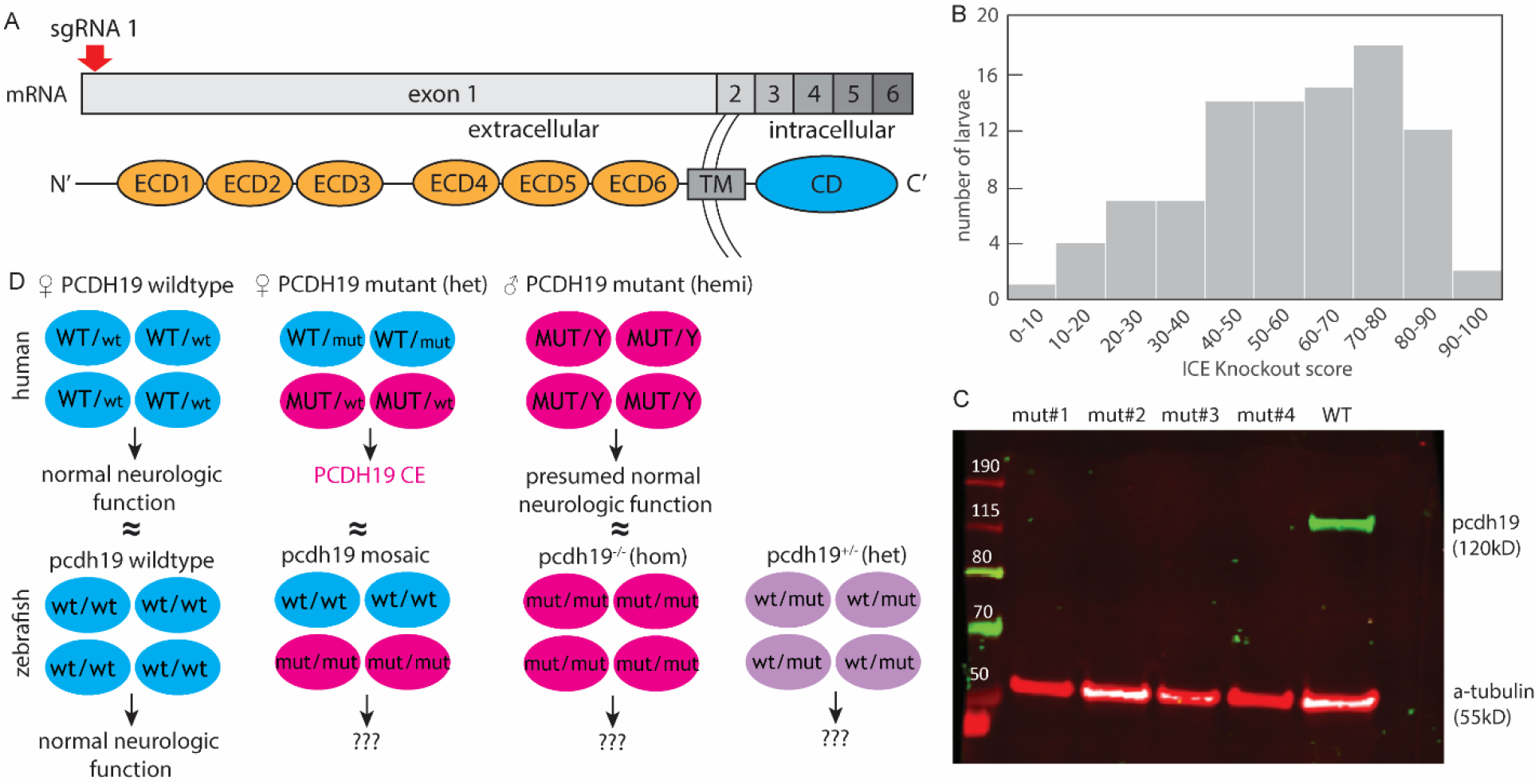
Mutant *pcdh19* loss-of-function zebrafish model. (A) Mutations introducing a 44-nucleotide insertion and 1-nucleotide deletion were introduced in the *pcdh19* gene at exon 1 (coding for the extracellular cadherin domains, ECD) (TM, transmembrane domain; CD, C-terminal domain), resulting in a frameshift. (B) Histogram of 192 F0 larvae injected with *pcdh19* sgRNA illustrating the distribution of the KO score, representing the proportion of cells predicted to result in the knockout due to the presence of a frameshift. (C) Western blot showing loss of full-length Pcdh19 protein in four different generated *pcdh19*^-/-^ mutant lines while WT larvae have intact Pcdh19 protein expression. (D) Schematic to illustrate the proposed disease mechanism in male and female humans and how we approached modeling them in zebrafish (wt, wildtype; mut, mutant).

To better understand the role of *PCDH19* in the developing brain and the impact cellular Pcdh19 mosaicism has in neurodevelopmental epilepsy, we developed mosaic and non-mosaic models of PCDH19 disorder in zebrafish by CRISPR/Cas9-mediated gene editing. Importantly, the mosaic mutants are comprised of some cells expressing mutant *pcdh19* and some wildtype, most closely approximating the mosaic expression that occurs in heterozygous female patients with PCDH19-CE. Because *pcdh19* is not present on a sex chromosome in zebrafish, we had the opportunity to also create mutants with germline heterozygous and homozygous frameshift mutations, predicted to lead to premature truncation and consequent loss of one or both copies of *pcdh19*. This provides the unique opportunity to evaluate the effects of mosaic and non-mosaic *pcdh19* variants in the zebrafish system, which is well-suited to evaluate features of epilepsy and neurodevelopmental dysfunction (Griffin et al., 2017). Given the human disease presentation of PCDH19-CE with seizures (spontaneous and fever-induced), learning defects, and other neuropsychiatric features, we assayed our models for zebrafish correlates of these features. We also performed neuroanatomic analyses to understand the mechanisms by which both mosaic and non-mosaic Pcdh19 loss-of-function (LOF) may lead to phenotypes in larval zebrafish.

## Results

### Generation of *pcdh19* zebrafish mutants

We generated zebrafish mutant lines with frameshift mutations that result in premature stop codons in exon 1 of the *pcdh19* gene (Figure 1A). We studied stable heterozygous (*pcdh19*^+/-^) and homozygous (*pcdh19*^-/-^) knockout (KO) zebrafish lines as well as mosaic F^0^ larvae that were acutely injected with the same guide RNA targeting exon 1 of *pcdh19*, which results in mosaic expression of Pcdh19 across all Pcdh19-expressing cells. Presence of the mosaic mutation was verified by DNA sequencing from *pcdh19* sgRNA-injected F^0^ larvae (after the experiments described below), and a ‘knockout score’ (reflecting the percentage of cells bearing the variant) was estimated by Interference of CRISPR Edits (ICE) analysis (Figure 1B). We observed highly variable numbers of indels generated by the CRISPR/Cas9 approach, which we leveraged in this study to mimic the tissue mosaicism present in females with *PCDH19* variants caused by random X-inactivation events during fetal development. Mosaic mutants with KO scores of >40 were used for subsequent experiments. We excluded the possibility that maternal wildtype mRNA is present early in development, either at the stage when we are evaluating larvae for phenotypes or before and exerting a lingering effect in our mosaicism model: we extracted mRNA from unfertilized eggs as well as embryos at 2hpf, 4hpf, and 24hpf and assessed the amount of *pcdh19* mRNA through quantitative PCR (qPCR). The levels of pcdh19 mRNA were undetectable in the unfertilized eggs and in the 2hpf and 4hpf embryos (Supp1.Fig. 1), indicating that negligible or no amount of maternal *pcdh19* mRNA is, likely to play a role in influencing the phenotypes we are evaluating during development. Successful loss of full-length Pcdh19 protein in different knockout mutants was verified by Western blot (Figure 1C). These different Pcdh19 LOF models reflect the human condition and give us the unique opportunity to study the effects of mosaic and non-mosaic Pcdh19 disruption (Figure 1D). None of our generated *pcdh19* mutant fish lines showed gross morphological abnormalities. Head and body size and shape were normal, general mobility, fertility, and survival (to at least 2 years) were intact.

### Increased spontaneous firing activity in zebrafish larvae with *pcdh19* mutations

To determine whether mosaic or non-mosaic *pcdh19* mutations result in abnormal neuronal network activity, we measured local field potentials (LFPs) in the optic tectum of 6-7dpf zebrafish larvae (Figure 2), a brain region known to have abnormal neuronal organization in the setting of Pcdh19 dysfunction (Cooper et al., 2015). In general, we distinguished between two types of spiking events that occurred in all our recordings at varying frequencies: events with a narrow peak and small amplitude (SAE) and events with a broad peak and large amplitude (LAE) (Figure 2A). LFP traces of wildtype larvae only occasionally captured the high-voltage events (approx. 0.2-0.4mV), while KO mutants displayed a range of epileptiform abnormalities. We often observed medium-to high-voltage deflections clustered in bursts and sometimes also in isolation. Discrete epileptiform discharges were 150-500ms long and resulted in 0.2-0.3mV sharp or blunt deflections. Quantification revealed a significantly increased LAE spike rate in mosaic and non-mosaic mutant *pcdh19* lines compared to their respective controls, while the three different control groups showed a uniformly low LAE spike rate (Figure 2B). LAE amplitude was significantly increased only in non-mosaic *pcdh19*^+/-^ KO larvae (Figure 2C). Quantification of burst frequency revealed abnormal clustering of large spiking events in the mosaic and non-mosaic *pcdh19*^+/-^ mutants compared to control larvae (Figure 2D). Similar but less frequent epileptiform abnormalities were observed in homozygous *pcdh19*^-/-^ KO zebrafish larvae. We confirmed these abnormal electrographic patterns in the form of increased and clustered bursts of abnormal neuronal firing in heterozygous and homozygous KO larvae in an independent *pcdh19* KO line with a frameshift mutation introduced at exon 2 (Suppl.Fig2A-D). These results indicate that a disruption of Pcdh19 function results in increased neuronal activity in *pcdh19* mutant fish, independent of mosaicism. Taken together, we found epileptiform activity in the form an increased neuronal spike rate in the tectum of mosaic and KO larvae that is clustered in bursts in mosaic and heterozygous KO larvae.

**Figure 2:**
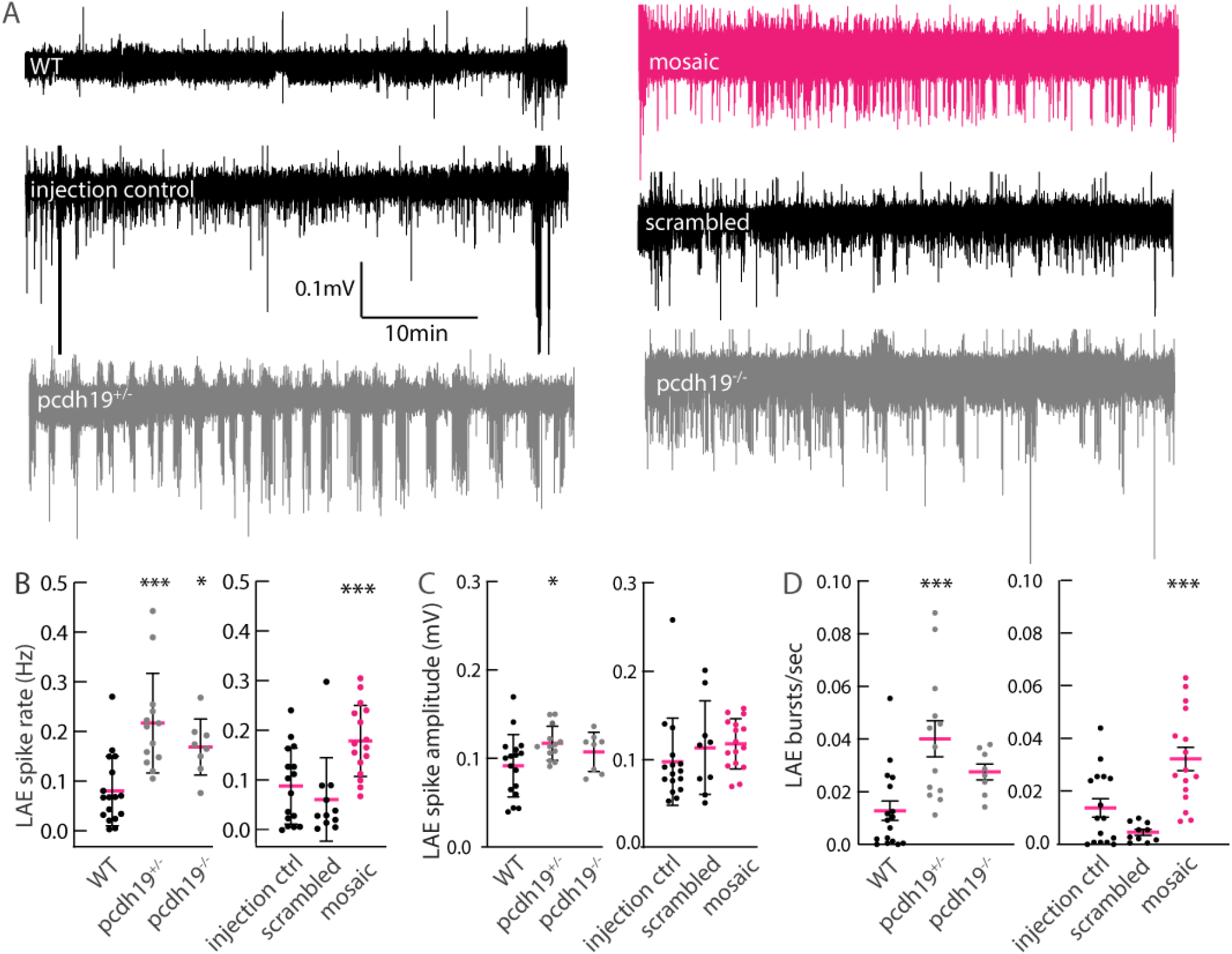
Hyperexcitability in LFP recordings of *pcdh19* mutant larval tectum. (A) Exemplary traces of LFP recordings for control and *pcdh19* mutants showing large amplitude events (LAE) consisting of bursts of spikes present in *pcdh19* mutants, which are rare in controls. (B) Quantification shows an increase in LAE rate in all *pcdh19* mutant lines, compared to their respective controls; mosaics were compared to scrambled injected and a ‘control injected’ group representing larvae that were injected with *pcdh19* sgRNA but had <10% indel frequencies. (C) The average LAE spike amplitude is increased in *pcdh19*^*+/-*^ KO mutant lines compared to WT controls. (D) The average amount of bursts is increased in *pcdh19*^*+/-*^ KO and mosaic mutants compared to their respective controls. n=16 WT, n=13 *pcdh19*^+/-^, n=8 *pcdh19*^-/-^, n=16 mosaic, n=15 injection control, n=11 scrambled injected. One-way ANOVA with Dunnett’s multiple comparisons test, *p<0.05, **p<0.01, ***p<0.001.

In complementary experiments, we further tested for hyperexcitability as a result of *pcdh19* dysfunction. First, we used the genetically encoded calcium sensor GCaMP6s to assess whole brain activity using lightsheet microscopy (Turrini et al., 2017). Cytosolic GCaMP6s is expressed pan-neuronally (elav3 promoter) and visualizes calcium influx during neuronal activity as changes in fluorescence (Figure 3A). We observed abnormal network activity in the tectum of a Pcdh19^+/-^ KO mutant, showing distinct clusters of neurons in the tectum with short bursts of increased activity (Figure 3B). We also observed longer-duration periods of dramatically increased whole brain activity (Figure 3C,D) lasting 1 to 2 min (Suppl. Video 1). Second, we used the neuronal activity marker phospho-ERK (pERK) to visualize and identify brain areas with increased neuronal activity 10 min prior to examination (Thyme et al., 2019). Whole brain antibody staining against pERK relative to total ERK (tERK) revealed increased pERK fluorescence signals within different parts of the tectum mainly in mosaic mutants and to a lesser extend in *pcdh19*^-/-^ mutants relative to control animals. While mosaic mutants primarily showed an increase in pERK signal in the neuropil, where neuronal processes converge, homozygous mutants had a decrease in parts of the medial tectum (Figure 3E); and heterozygous mutants showed no difference vs. controls. To corroborate that the increased brain activity in mosaic fish can be attributed to Pcdh19 dysfunction, we generated mosaic *pcdh19* mutants using a different sgRNA targeting *pcdh19* at exon 2. We observed a similar pattern of increased pERK signal in the tecum of these larvae (Supp2.Fig1E). These experiments indicate that abnormal brain activity might be limited to the optic tectum in larval zebrafish. Together, these results provide further evidence that Pcdh19 dysfunction results in altered brain activity. However, using pERK as a marker for neuronal activity only showed a significant increase in the tectum of mosaic mutants.

**Figure 3:**
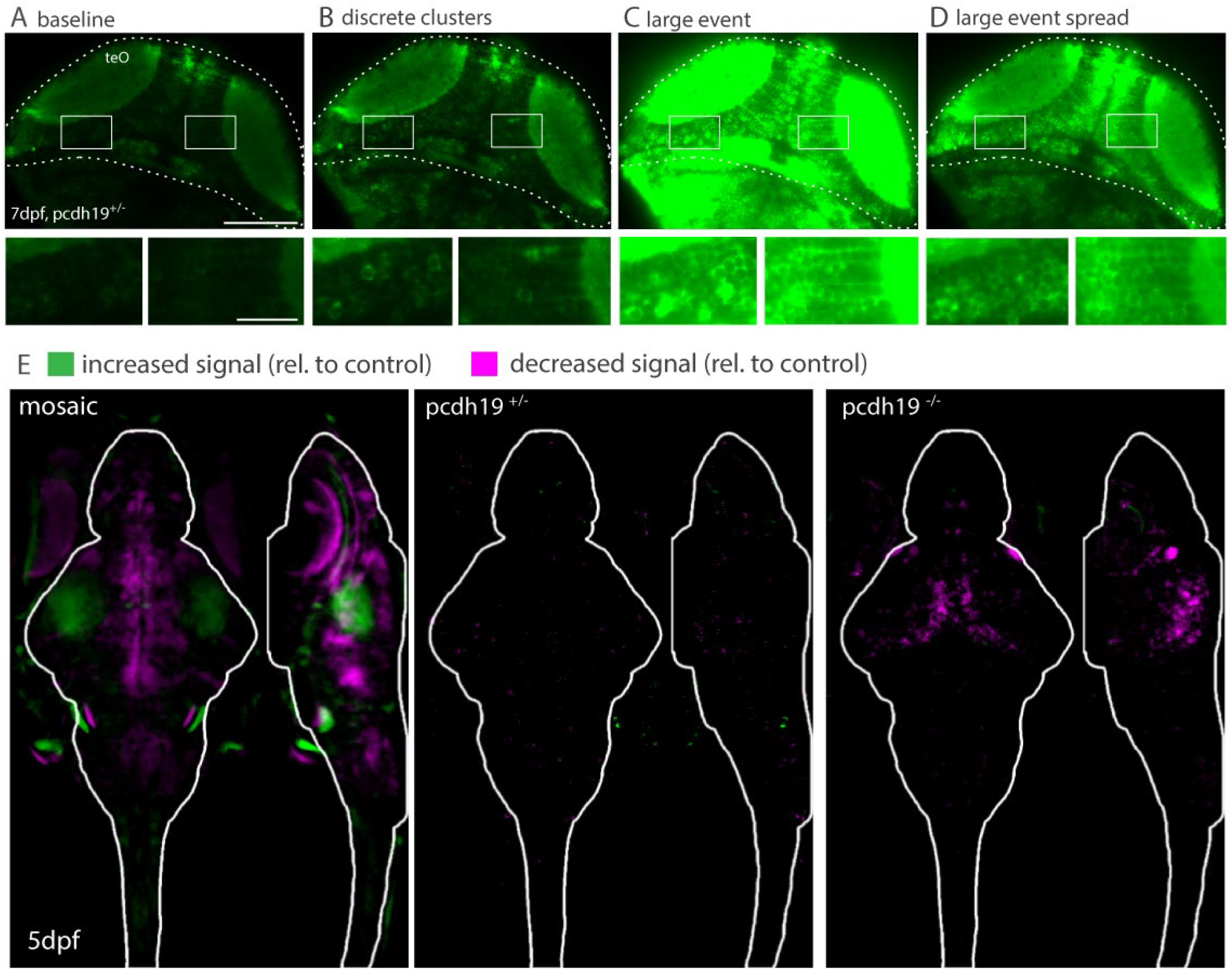
Hyperexcitability in *pcdh19* mutants. (A) Lightsheet images of a GCaMP6s-expressing *pcdh19*^+/-^ mutant fish at baseline. Scale bar 100µm and 20µm in higher magnification images below. teO; tectum opticum. (B) Increased GCaMP fluorescence in two different neuronal clusters of the tectum. (C) Dramatic increase in GCaMP fluorescence across the entire brain of a *pcdh19*^+/-^ mutant fish during a large discharge event. (D) Slowly attenuating GCaMP intensity after the large event. (E) Sum of slices projection of confocal images show pERK signal intensity is increased in retinotectal processes in mosaic *pcdh19* mutant larvae, no change in *pcdh19*^+/-^ mutants, and decreased signal in *pcdh19*^-/-^ mutants, n=18 mosaic, n=28 scrambled injected, n=82 *pcdh19*^+/-^ and n=28 *pcdh19*^-/-^ larvae.

### Mosaic and non-mosaic mutations in *pcdh19* do not lead to a consistent seizure phenotype

To assess whether the abnormal patterns of electrical activity observed in the LFP recordings and in calcium imaging correlate with seizure-like activity, we subjected mutant and control larvae to high-throughput assays to further characterize the range of abnormalities resulting from Pcdh19 dysfunction. We evaluated for behavioral seizure-like events in zebrafish larvae, defined as short episodes of hyperlocomotion that result in rapid swimming (Baraban et al., 2013) that can either occur spontaneously or can be evoked by proconvulsant compounds or temperature elevation (hereafter referred to as seizures). At baseline, we observed only occasional seizures in WTs (2.5%), which was comparable to the *pcdh19*^+/-^ (2%), *pcdh19*^-/-^ (0.7%), and mosaic (0.3%) groups (Figure 4A). When challenged with a concentration of PTZ (0.1 mM) that is not sufficient to generate seizures in WT larvae (a ‘subthreshold’ concentration), a significantly higher proportion of *pcdh19*^+/-^ mutant larvae exhibited seizures than WT larvae (4.2% vs 0.9% WT) (Figure 4A). Seizing *pcdh19*^+/-^ mutants also had more seizures over the duration of the trial than the WT larvae (Figure 4A). Slowly increasing the bath temperature, and consequently larval body temperature, from 22°C to 37°C resulted in a few seizures in WT larvae (<2%) and no significant changes from the baseline seizure frequency in the *pcdh19* mutant groups: *pcdh19*^+/-^ (3.1%), *pcdh19*^-/-^ (1.2%), mosaic *pcdh19* mutants (4.6%) (Figure 4A). Increasing the PTZ concentration to 2.5 mM, a concentration at which most animals are expected to develop seizures, significantly more *pcdh19*^+/-^ KO larvae (99.5%) had at least one seizure compared to WT larvae (90.8%) (Figure 4B). The average number of seizures in each group was similar (13.1% seizures/WT vs 13.7% seizures/*pcdh19*^+/-^ larvae) (Figure 4B). In some forms of epilepsy, seizures can be triggered by intermittent photic stimulation; we therefore challenged mutant larvae with exposure to different light-dark paradigms and strobed frequencies. Although the *pcdh19*^+/-^ and *pcdh19*^-/-^ KO mutants moved on average more and with higher velocities during dark periods, we did not detect striking differences or seizures in response to different strobe light frequencies between groups (Figure 4D). To summarize, only the non-mosaic heterozygous *pcdh19* KO mutants showed a slightly increased susceptibility to chemically provoked seizures while the mosaic and non-mosaic *pcdh19*^*-/-*^ KO mutant larvae did not have an increase in seizure events compared to their controls in any of the tested assays. We did not observe a correlation between the electrographic abnormalities observed during LFP recordings in the mosaic and KO mutant lines and the occurrence of gross behavioral seizures.

**Figure 4:**
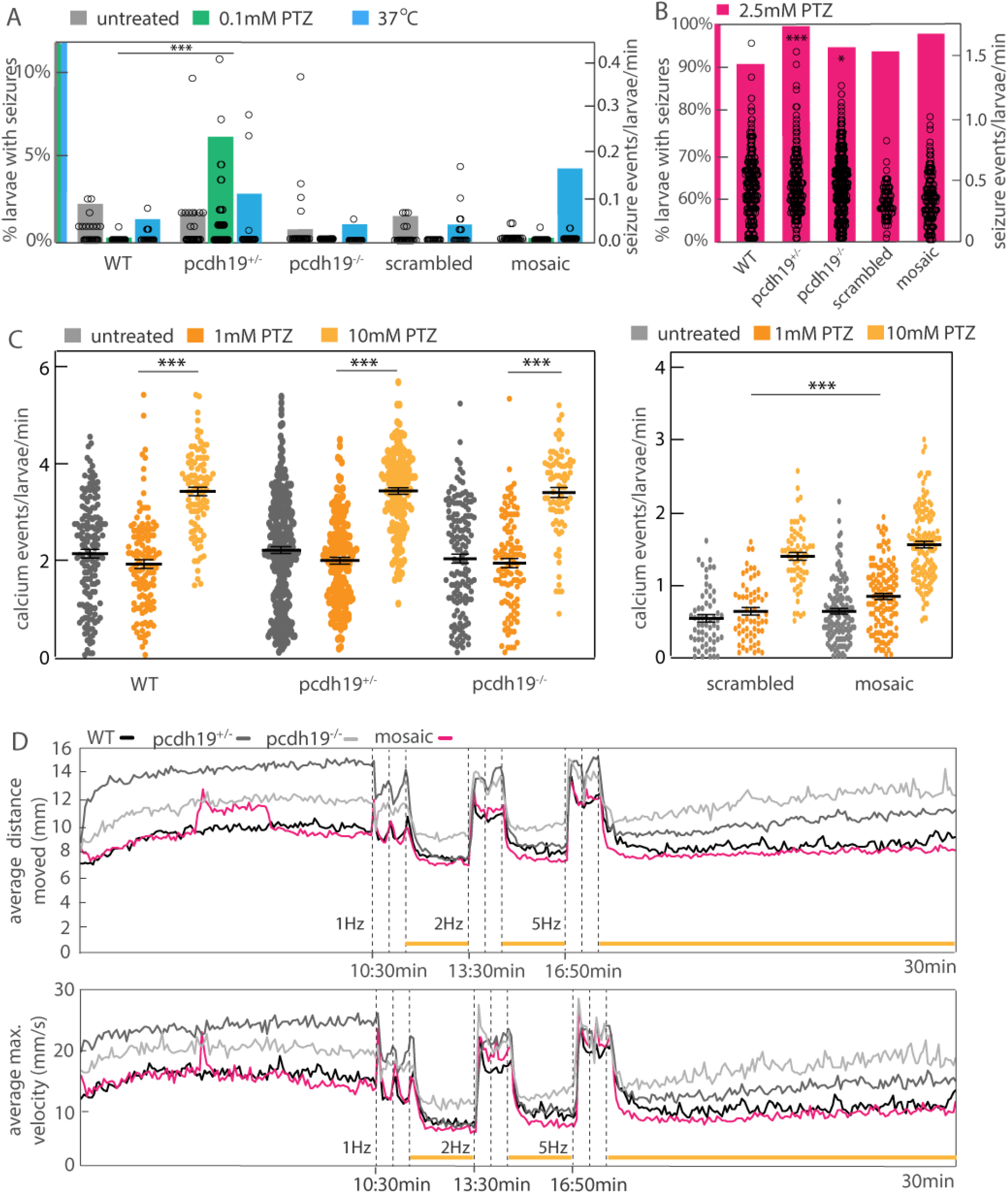
High-throughput detection of spontaneous and provoked seizure activity by behavioral and calcium fluorescence-based assays. (A) Quantification of the percentage of larvae with seizure events (left axis, green bar=0.1mM PTZ exposure blue bar=exposure to 37°C warm water) and the number of seizures per larvae per minute (right axis, open circles) untreated (grey bars) or after exposure to 0.1mM PTZ (green bars) or temperature increase up to 37°C (blue bars). *pcdh19*^+/-^ larvae treated with 0.1mM PTZ showed an increased number of seizures compared WT. Untreated and 0.1mM PTZ: n=429 WT, n=500 *pcdh19*^+/-^, n=539 *pcdh19*^-/-^, n=384 mosaic, n=192 scrambled, Chi-squared test, ***p<0.001. Untreated and temperature elevation: n=258 WT, n=96 *pcdh19*^+/-^, n=163 *pcdh19*^-/-^, n=150 mosaic, n=192 scrambled, Chi-squared test, *p<0.05, ***p<0.001. (B) Quantification of seizure events in larvae treated with 2.5mM PTZ (left axis, pink bars=2.5mM PTZ) induces seizures in almost all *pcdh19*^+/-^ and pcdh19^-/-^ mutant larvae, significantly more than in WT larvae. n=447 WT, n=225 *pcdh19*^+/-^, n=620 *pcdh19*^-/-^, n=192 mosaic, n=192 scrambled, Chi-squared test, *p<0.05, ***p<0.001. The frequency of seizures in each animal is similar across genotypes (right axis, open circles) (C) Quantification of the average event rate of calcium transients in GCaMP6s larvae shows no difference between WT and mutants in the untreated condition; there is a significant increase in events in all groups after exposure to 10mM PTZ, similar to WT. Comparing mosaic and scrambled injected larvae shows a significant increase in calcium events when treated with 1mM PTZ. n=155 WT, n=264 *pcdh19*^+/-^, n=145 *pcdh19*^-/-^, n=128 mosaic, n=61 scrambled, one-way ANOVA with Tukey’s multiple comparisons test for Pcdh19 KO mutants and two-way ANOVA with Sidak’s multiple comparisons test mosaic vs scrambled and PTZ vs no treatment, *p<0.05, ***p<0.001. (D) Average movement (distance moved in top panel and maximum velocity in lower panel) of larvae exposed to dark, 100% light (indicated by horizontal yellow lines), or 3 pulses, 3 sec each, of strobe light (1Hz, 2Hz, and 5Hz, indicated by vertical dashed lines) over 30 min did not change in response to the different light paradigms in *pcdh19* mutants compared to WT. n=223 WT, n=227 *pcdh19*^+/-^, n=134 *pcdh19*^-/-^, n=261 mosaic.

Since our behavioral seizure detection and analysis might not be sensitive enough as a readout to capture the hyperexcitability phenotype of *pcdh19* mutants, we performed low-resolution, high-throughput calcium imaging to assess for and characterize seizure-like activity. We observed *pcdh19*^+/-^ and *pcdh19*^-/-^ KO larvae with stereotypical rapid, swirling swim behaviors and a simultaneous increase in GFP fluorescence, likely representing seizures with increased brain activity that subsided after the events (Suppl. Video 2, 3). However, quantification with our event detection algorithm did not capture a difference in the overall rate of high calcium events between untreated *pcdh19* KO mutants and WT or scrambled injected vs mosaic animals (Figure 4C).

Interestingly, PTZ treatment provoked on average slightly more calcium events in mosaic *pcdh19* mutants compared to scrambled injected controls (Figure 4C). However, again we did not observe a clear correlation between the neuronal hyperexcitability that occurs in mosaic an KO mutants and an increase in high calcium spiking events in this experimental setting since there was no consistent increase in neuronal calcium activity in mosaic and KO mutant pcdh19 larvae compared to control larvae. These results suggest that low-resolution microscopy is not suited to to visualize changes in neuronal firing patterns that occur in all *pcdh19*^+/-^ mutant lines, given the low number of synchronously firing neurons within a small brain area, as suggested in Figure 3B.

### Fewer inhibitory neurons in the tectum of *pcdh19*^+/-^ mutant larvae

To probe for a structural defect that might explain the abnormal neuronal excitability in *pcdh19* mutant larvae in the transgenic background Tg(dlx6a-1.4kbdlx5a/dlx6a:GFP::vGlut:DsRed), we aimed to determine the number of inhibitory and excitatory neurons present in mutants vs. WT conditions. At 5dpf, WT transgenic larvae express GFP in distinct neuronal populations clustering in the tectum, forebrain, and midbrain, while dsRed expression is found in all brain areas with large clusters within the olfactory globes, midbrain, and parts of the hind brain (Figure 5A). Across the whole brain, no gross differences in the expression patterns and distribution of excitatory and inhibitory neurons were observed across all *pcdh19* mutant larvae. However, regional quantification revealed a difference in the number of inhibitory neurons within the tectum (Figure 5B). On average, fewer inhibitory neurons were present in the tectum of 5dpf *pcdh19*^+/-^ KO zebrafish larvae compared to age-matched control larvae or *pcdh19*^-/-^ KO mutants. Comparison of scrambled injected vs. mosaic *pcdh19* mutant larvae showed no difference in the number of inhibitory neurons in the tectum, suggesting that only the non-mosaic *pcdh19*^+/-^ KO fish display this interneuron phenotype, correlating with the observation that these mutants also have the most pronounced behavioral hyperexcitability phenotype (as seen in Fig.4).

**Figure 5:**
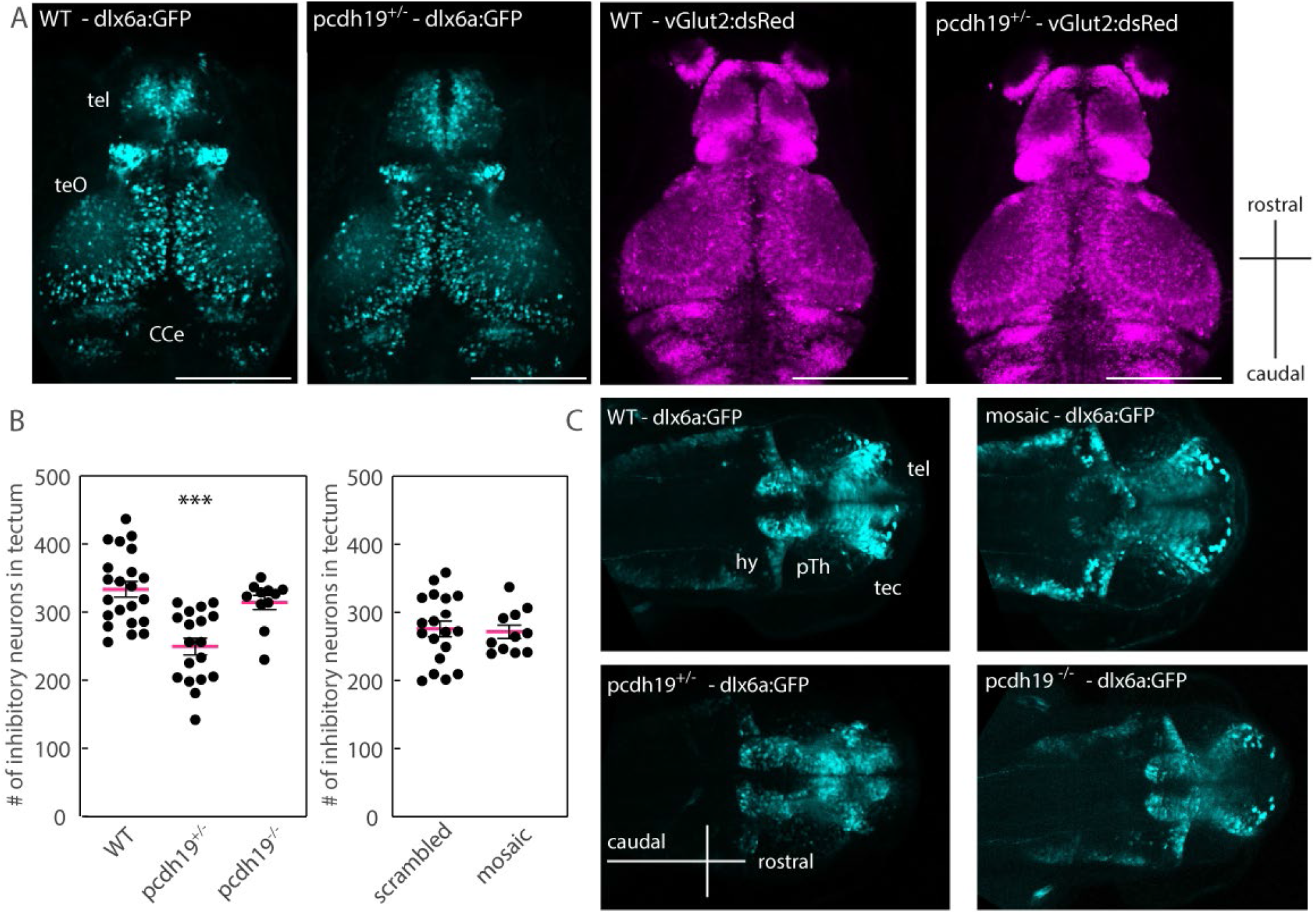
Brain structural abnormalities in *pcdh19*^+/-^ larvae. (A) Representative confocal maximum intensity projections of a Z-stack taken from a 5dpf WT and a *pcdh19*^+/-^ mutant crossed with the Tg(dlx6a-1.4kbdlx5a/dlx6a:GFP::vglut2:DsRed) line show no gross morphological abnormalities in either the dsRed or the GFP expressing neuron population. Scale bar 200µm. (B) Quantification of the number of inhibitory neurons within the tectum shows a reduction only in *pcdh19*^+/-^ mutants compared to WT controls. n=22 WT, n=19 *pcdh19*^+/-^, n=11 *pcdh19*^-/-^, n=19 mosaic, n=11 scrambled larvae, one-way ANOVA with Dunnett’s multiple comparisons test, p<0.0001. (C) Representative confocal maximum intensity projections of a Z-stack of 2dpf WT, *pcdh19*^+/-^, *pcdh19*^-/-^ and mosaic mutants showing an altered, immature neuronal pattern in the *pcdh19*^+/-^ mutant compared to the WT larvae. tel, telencephalon; teO, tectum opticum; CCe, cerebellar corpus; tec, tectum; pTh, pre-thalamus, hy, hypothalamus.

To determine whether the reduced total number of inhibitory neurons, noted at 5dpf, in these mutants is due to increased cell death, we performed TUNEL assays at 2, 3 4, and 5dpf in *pcdh19* KO mutants. Quantification did not show increased cell death in any of the lines at any investigated time point, suggesting that destruction of developing interneurons is not the mechanism for the observed interneuron phenotype (Suppl. Fig. 3). Quantification of the number of inhibitory neurons at 2dpf was impossible since migrating cells form dense clusters and thus, we were unable to identify single GFP-expressing cells at this age. However, we observed differences in the developing brain of *pcdh19*^+/-^ larvae. While inhibitory neurons in WT embryos start to form distinct clusters in the thalamic brain regions at 2dpf, it appeared that *pcdh19*^+/-^ mutant brains had immature patterns of GFP-expressing neurons with overall smaller cell clusters (Figure 5C). These are reminiscent of earlier stages of brain development, suggesting transiently delayed development that recovers to normal by 5dpf. Quantification showed a significantly reduced GFP-fluorescent area in *pcdh19* KO larvae compared to WT larvae, indicating either that fewer inhibitory neurons are present, or they are more densely clustered at this stage (Suppl. Fig 2F). Together these results suggest that Pcdh19 plays a part in the development of the inhibitory neuronal network and a disruption of Pcdh19 function appears to have a stronger effect in heterozygous *pcdh19* KO animals than mosaic mutants. The observation of reduced inhibitory neurons only in heterozygous KO mutants is in line with the overall strongest behavioral seizure phenotype in heterozygous KO mutants (as seen in Fig.4A and B).

### Mutant *pcdh19* zebrafish do not display evidence of anxiety or impaired spatial learning

Because patients with *PCDH19* variants often display neuropsychiatric comorbidities with intellectual disability and/or autism, behavioral dysregulation, and obsessive features (Samanta, 2020), we evaluated our *pcdh19* zebrafish mutants for learning defects and features of anxiety, which have established correlates in zebrafish. We found no difference between mutant and control larvae using the thigmotaxis, or wall-hugging, assay for larval zebrafish, which is considered a correlate for anxiety (Schnorr et al., 2012). WT and scrambled injected larvae spend on average 71.8% and 70.9% of the time ‘hugging’ the wall, while this proportion is increased to 88.5% in WT larvae treated with the anxiogenic caffeine, indicative for increased anxiety levels (50 mg/L) (Figure 6A and B). We did not observe a statistically significant difference in the wall-hugging behavior of *pcdh19*^*-/-*^ KO or mosaic mutants compared to their respective controls, and we observed a significant reduction of thigmotaxis in *pcdh19*^+/-^ fish. As neuropsychiatric dysfunction tends to exacerbate with age in human PCDH19-CE, we also looked at older *pcdh19* mutant fish. We found weak evidence for anxiety-like behavior only in *pcdh19* mosaic mutants in the bottom dwelling test using adult zebrafish (Figure 6C,D,E). While WT fish spent on average 93±61sec in total in the top compartment of the test tank, *pcdh19*^+/-^ and *pcdh19*^-/-^ fish tend to spend more of time in the top (127±85, 151±56sec respectively), indicating no anxiety-like behavior. Scrambled and mosaic injected adults spent even more time in the top compartment but to a similar degree (Figure 6D). Interestingly, the mosaic *pcdh19* mutants had significantly more zone transitions compared to the scrambled injected and *pcdh19* KO fish (Figure 6E), a behavior that is consistent with anxiety-like behavior. However, this group was also the only group in which almost all of tested animals entered the top compartment at least once, while in all other groups a substantial proportion of fish remained motionless at the bottom of the tank (Figure 6F). Based on these findings, we conclude that *pcdh19* KO mutant zebrafish do not display a strikingly increased anxiety-like phenotype, while *pcdh19* mosaic mutants show tendencies towards an anxiety-related phenotype.

**Figure 6:**
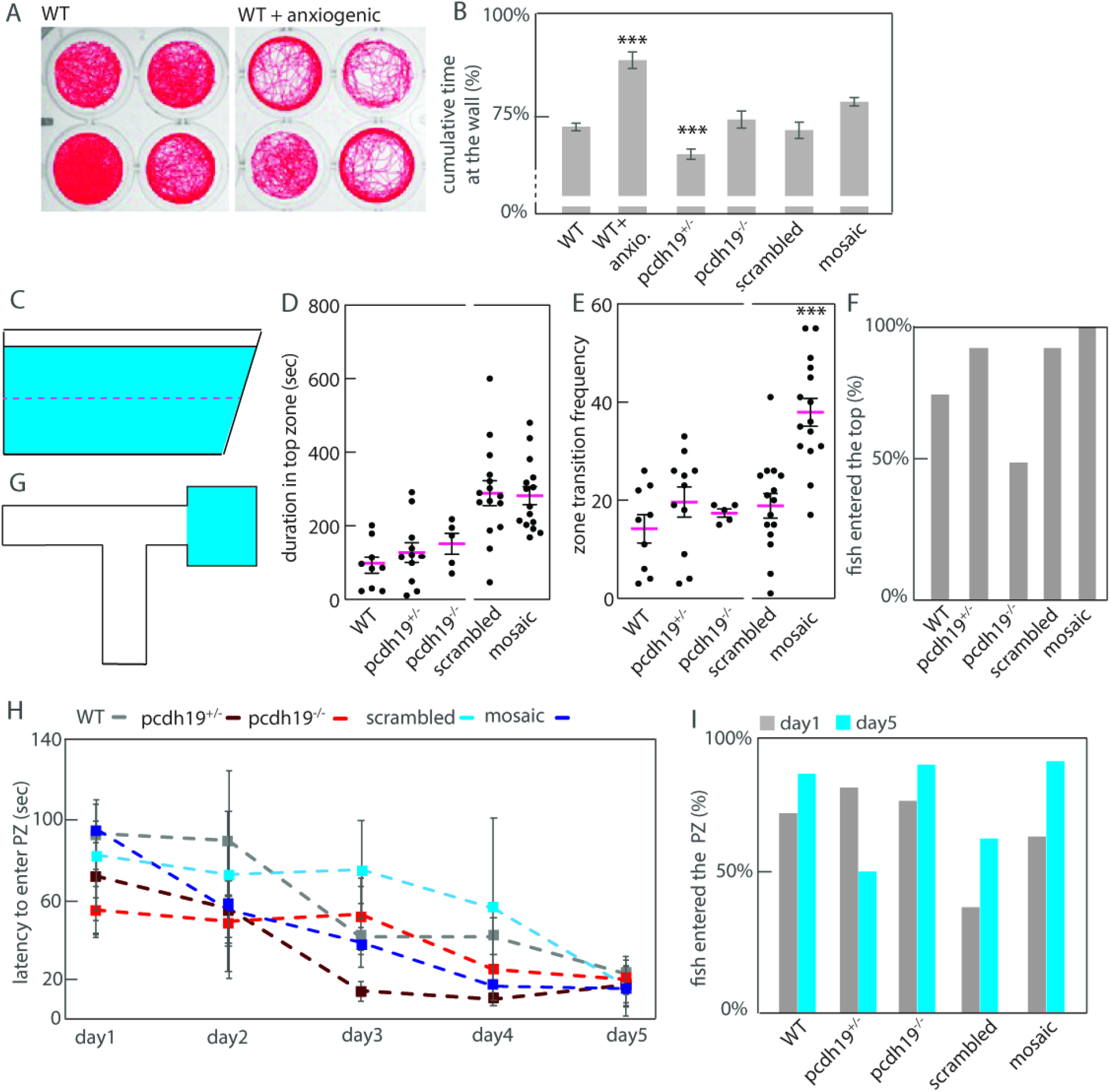
Behavioral paradigms to assess for features of anxiety and learning defects in zebrafish. (A) Movement plots of 4 representative WT larvae untreated and treated with the anxiogenic caffeine in a 24-well plate over 30 min. (B) Quantification of the time spent at the wall shows only larvae treated with the anxiogenic spend more time at the borders of the well, while *pcdh19*^+/-^ larvae spent significantly less time close to the wall. n=367 WT, n=311 *pcdh19*^+/-^, n=91 *pcdh19*^-/-^, n=283 mosaic, n=60 scrambled, n=59 WT+caffeine treated leave, one-way ANOVA with Tukey’s multiple comparisons test, ***p<0.001. (C) Schematic of the bottom-dwelling test tank. (D) Quantification of the duration the adult zebrafish spent in the top compartment of the test tank, that is unchanged between control and *pcdh19* mutant animals. (E) The frequency of transitions between bottom and top compartment is significantly increased in mosaic *pcdh19* mutants compared to their controls. (F) The percentage of fish that entered the top compartment is lower for WT and *pcdh19*^-/-^ fish compared to the other groups. n=12 WT, n=12 *pcdh19*^+/-^, n=10 *pcdh19*^-/-^, n=15 mosaic, n=15 scrambled fish, one-way ANOVA with Sidak’s multiple comparisons test, p<0.0001. (G) Schematic of the T-maze tank. The blue compartment depicts the preference zone (PZ). (H) Average latency to enter the PZ on the 5 consecutive training days is similar for all compared groups. (I) The percentage of fish that entered the PZ on day1 or 5 varies between groups. n=36 WT, n=28 *pcdh19*^+/-^, n=36 *pcdh19*^-/-^, n=64 mosaic, n=16 scrambled fish, one-way ANOVA with Dunnett’s multiple comparisons test.

Using the T-maze test with adult zebrafish (Figure 6G), we found no defects in the learning capabilities of *pcdh19* mutants. Both, the performance of mutant and the WT increased similarly over the duration of the trial. The average latency for WT zebrafish to reach the preference zone (PZ) went down from 93.7±76s on day 1 to 23.5±41s on day 5, this was a 4fold and statistically significant improvement in reaching the preference zone (Figure 6H). Comparably, all mutant *pcdh19* zebrafish improved significantly from day1 to day5 with mosaic and *pcdh19*^+/-^ animals showing the most dramatic response (with 6.4-and 4.1-fold improvement respectively), *pcdh19*^-/-^ performance improved the least (2.7-fold improvement relative to day 1). Comparing latency to reach the preference zone on day 1 or on day 5 between WT and *pcdh19* mutants did not show any significant differences (Figure 6H). However, a considerable number of fish were excluded from the analysis because they either failed to move during the 5min recording period or failed to reach the preference zone (Figure 6I). Together, these results show that both, mosaic and non-mosaic *pcdh19* mutants did not show overt signs of increased anxiety or cognitive decline compared to age-matched control animals, indicating that distinct Pcdh19 expression patterns have no severe impact on learning and memory or anxiety in 4-to 5-month-old zebrafish.

## Discussion

A prevailing hypothesis is that human *PCDH19*-associated disease phenotypes result from the presence of cellular mosaicism in the brain, with clusters of neurons expressing WT and others expressing mutant PCDH19 protein. To test this hypothesis, we assayed both mosaic and non-mosaic *pcdh19* loss-of-function zebrafish larvae for features related to epilepsy and comorbid features associated with human disease, as well as for underlying neurodevelopmental abnormalities that might be responsible for these features. Our most striking finding was abnormal hyperexcitability in mosaic and non-mosaic LOF larvae, as evidenced by spontaneous epileptiform discharges observed by electrophysiological recording. This is a feature associated with epilepsy previously shown in a zebrafish model for Dravet syndrome (Baraban et al., 2013). These findings were confirmed by using multiple *pcdh19* LOF lines and by acutely injecting a different sgRNA to target *pcdh19* at a distinct genomic position. Surprisingly, we saw only limited indications of abnormal excitability using GCaMP-mediated calcium imaging and behavioral based locomotor activity paradigms both at baseline and during provoking conditions using light, heat and the proconvulsant PTZ at different concentrations, all established methods to assay hyperexcitable states in zebrafish (Afrikanova et al., 2013; Hortopan et al., 2010; Hunt et al., 2012; Liu and Baraban, 2019). Using these assays, the strongest hyperexcitability phenotype was observed in non-mosaic, heterozygous *pcdh19* mutant larvae showing spontaneous and PTZ-induced behavioral seizures as well as abnormal calcium events. We conclude that the nature of the pathology, and its resultant hyperexcitability, generated by mosaic and non-mosaic *pcdh19* LOF mutations in zebrafish larvae requires a sensitive detection method (tectal LFP recordings) to consistently identify evidence of hyperexcitability in all lines. Indeed, the lack of seizure-like events using behavioral assays in the setting of a hyperexcitable neuronal network is not an uncommon phenomenon, as recently demonstrated in a large zebrafish epilepsy screen (Griffin et al., 2021).

Relative changes in pERK antibody fluorescence, employed to provide neuroanatomical localization of neuronal activity (Thyme et al., 2019), showed complex results in our mosaic and non-mosaic *pcdh19* mutants. Mosaic mutants showed marked changes in neuronal activity in a widespread pattern which partly overlaps with tectum, but heterozygous *pcdh19*^*+/-*^ mutants—the same genotype that showed evidence of hyperexcitability by other modalities—showed more subtle abnormalities using pERK fluorescence. Since this method only visualizes changes in neuronal activity that occurred approximately 10min prior to preparation, it may not be ideally suited to image short, transient events such as seizures. The abnormal pERK patterns found in the mosaic mutants may therefore represent longer lasting, persistent changes in neuronal activity patterns that are different from the non-mosaic *pcdh19* lines and may not represent the presence of seizures as such but rather a baseline, tonic hyperexcitable state.

In order to further explain the *pcdh19* mutant hyperexcitability phenotype, we assessed neuronal numbers. We observed reduced numbers of inhibitory neurons at 2dpf and 5dpf with normal TUNEL staining in heterozygous LOF mutants. This observation represents a change that may underlie with a hyperexcitable neuronal network. Our results suggest a failure of development of inhibitory neurons in heterozygous *pcdh19 KO* mutants that is not observed in mosaic LOF mutants, Interestingly this correlates with the more severe hyperexcitability phenotype present in heterozygous *pcdh19* mutants. Consistent with our findings, experiments silencing Pcdh19 in cultured rat neurons showed reduced GABAaR and GAD65/67-positive puncta and reduced mIPSC frequency with slower decay kinetics, suggesting a *Pcdh19*-mediated mechanism is necessary for the composition and function of the inhibitory synapse (Bassani et al., 2018).

Common neuropsychiatric features of the human PCDH19-related disorder include intellectual disability, autism, and obsessive-compulsive disorder which tend to worsen over time (Samanta, 2020). We therefore evaluated not only larval but also adult zebrafish for these abnormalities. Similar to published reports in mice that assessed changes in anxiety and memory-related behavior (Hayashi et al., 2017; Pederick et al., 2016), none of the adult fish displayed consistent or striking abnormalities in behavior using two different tests that assess cognitive performance and anxiety-like states. The discrepancy between the severe neuropsychiatric comorbidities in humans and the rather mild reported phenotypes in mice and zebrafish, if any, suggest that human brain function is more sensitive to PCDH19 LOF or that existing animal models are less sensitive either due to genetic background effects, limitations of existing behavioral assays to reveal pcdh19-related phenotypes, or a unique requirement for PCDH19 in the human brain.

The extent to which non-mosaic Pcdh19 LOF contributes to abnormal hyperexcitability and epilepsy remains an unresolved question. As the prevailing theory postulates that cellular interference in the setting of mosaic LOF is required for a phenotype (Pederick et al., 2018), we conclude that an alternative mechanism based on haploinsufficiency must play a role to account for our observations that are consistent with hyperexcitability in non-mosaic heterozygous or homozygous *pcdh19* mutant zebrafish. This hypothesis is supported by the presence of behavioral abnormalities in human male carriers (our own unpublished observations), abnormalities in neuronal network development and function in the optic tectum of Pcdh19 mutant zebrafish (Light et al 2019; Cooper 2015), and abnormalities in *pcdh19* KO mice. This study showed heightened evoked seizure susceptibility in both female *Pcdh19*^+/-^ (X-linked mosaic) and female *Pcdh19*^-/-^ (non-mosaic) mice compared to WT and male hemizygous mice (Rakotomamonjy et al., 2020), strengthening the hypothesis that mosaicism-independent mechanisms also contribute to *PCDH19* disease phenotypes.

The differences and the lack of a clear correlation between genotype and phenotype that is seen across zebrafish with mosaic, heterozygous, and homozygous *pcdh19* mutations are reminiscent of differences observed in studies of other model systems, including mouse models of epilepsy (Wang and Frankel, 2021). Given that the protocadherin gene family is well-conserved across species, and that the zebrafish *pcdh19* gene shares 73% homology with human *PCDH19* (Blevins et al., 2011; Wu, 2005), the zebrafish systems allow for robust evaluation of phenotypes that correlate with human patients with pathogenic variants in *PCDH19*. Indeed, zebrafish models of other genetic epilepsies have been well described and even used successfully in drug screens (Baraban et al., 2013; Eimon et al., 2018; Grone et al., 2016). The lack of spontaneous behavioral seizures in the mosaic or non-mosaic Pcdh19 LOF fish, despite abnormal electrophysiological findings, does not diminish the significance of the positive findings observed and is consistent with several previously reported models of genes responsible for human epilepsy. A recent large-scale epilepsy screen reported a low prevalence of behavioral seizures in a set of 40 zebrafish lines with variants in catastrophic childhood epilepsy genes as well as surprisingly low prevalence of epileptiform abnormalities in electrophysiological recordings (Griffin et al., 2021). Due to technical limitations, we could not test for the presence of segregated and abnormally clustered cell populations in the brain in the mosaic mutants. Furthermore, zebrafish larvae are not sexually dimorphic and thus, this disease model does not allow for the evaluation of sex-dependent differences. Given that the most consistent phenotypic abnormality relied on electrophysiological experiments, we advocate for including such experiments in drug screens for the identification of novel therapeutic compounds to target pcdh19 dysfunction, which are readily feasible in zebrafish larvae (Griffin et al., 2021). Evaluation of some of these aspects in mouse models may allow additional understanding of the role of PCDH19 in brain development and epilepsy (Hoshina et al., 2021; Pederick et al., 2018).

In summary, we demonstrate that mosaic and non-mosaic *pcdh19* mutant zebrafish recapitulate several features of human PCDH19-CE, including the core feature of brain hyperexcitability in the zebrafish models that correlates with the network dysfunction resulting in epilepsy in human patients. We provide evidence suggesting that the mechanism for hyperexcitability in non-mosaic Pcdh19 mutants may involve abnormal inhibitory neuron development. Understanding the molecular basis of brain dysfunction that arises from mosaic and non-mosaic mutant Pcdh19 expression provides a fundamental step towards understanding PCDH19-CE disease mechanisms, fostering future studies to develop mechanism-based strategies to treat symptoms. Specifically, single-cell RNA sequencing experiments would help to identify abnormal molecular pathways in mosaic or KO loss of function mutants that could lay the foundation for targeted treatment options. For example, the reduction of inhibitory neurons during development suggests that modifying GABAergic activity, which can be achieved pharmacologically in patients with some classes of anti-seizure medications (e.g., benzodiazepines, barbiturates), may represent a more effective treatment mechanism than many of the other mechanisms of conventional anti-seizure medications.

## Material and Methods

### Zebrafish Maintenance

All zebrafish experiments were approved and conducted in accordance with the standards of the Boston Children’s Hospital (BCH) Institutional Animal Care and Use Committee (IACUC). Fish were maintained at 79°F on a 10h-14h dark-light cycle in a dedicated zebrafish core facility. All experiments were performed in *Danio rerio* strain AB or TAB.

### Guide RNA design and microinjections

Single guide RNAs (sgRNAs) were designed using the CHOPCHOP online tool v1 (https://chopchop.rc.fas.harvard.edu) and selected using its ranking algorithm based on target specificity and minimization of off-target effects. We used *pcdh19* sgRNAs targeting the beginning of exon 1 and exon 2 to maximize loss-of-function of the protein, based on our observation that most patient mutations are located on exon 1, with a few in exon 2, and almost none in the remaining exons. Microinjections into embryos at the one-cell stage were performed as described previously (Dentici et al., 2021) with a mixture of 1μl synthetic guide RNAs (*pcdh19* oligo 2/exon1: GGTGTATTTCAAATTGAACA or *pcdh19* oligo 3/exon 2: GGAGACGGACAAGATGAATG at 2000ng/μl, Synthego, Redwood City, CA, USA) and 1μl recombinant Cas9 protein (1 μg/μl, PNA Bio, Thousand Oaks, CA, USA) and 2μl RNAse-free water. For control experiments, a scrambled sgRNA (ACAAGGAGGTAGGCGAGAAC) that does not target zebrafish DNA was injected. A total of 2nl mix per embryo was injected using clipped glass capillaries and a microinjector.

### Genotyping and PCR

To determine sgRNA efficiency and to confirm the different genotypes in *pcdh19* KO fish, fin clips were cut from adult 3-month-old fish and DNA extracted using the HotShot protocol (Meeker et al., 2007). Euthanized larvae were dissolved in 50μl 50mM NaOH by heating for 20min to 95°C followed by addition of 5μl 1M Tris solution. Subsequent PCR using 2xPlatinum Taq polymerase and Sanger sequencing was performed with primers summarized in Table 1. The amount of indels in *pcdh19* mosaic larvae and the potentially resulting knockdown percentage was determined via Inference of CRISPR Edits (ICE) analysis (https://www.synthego.com/products/bioinformatics/crispr-analysis).

**Table 1:**
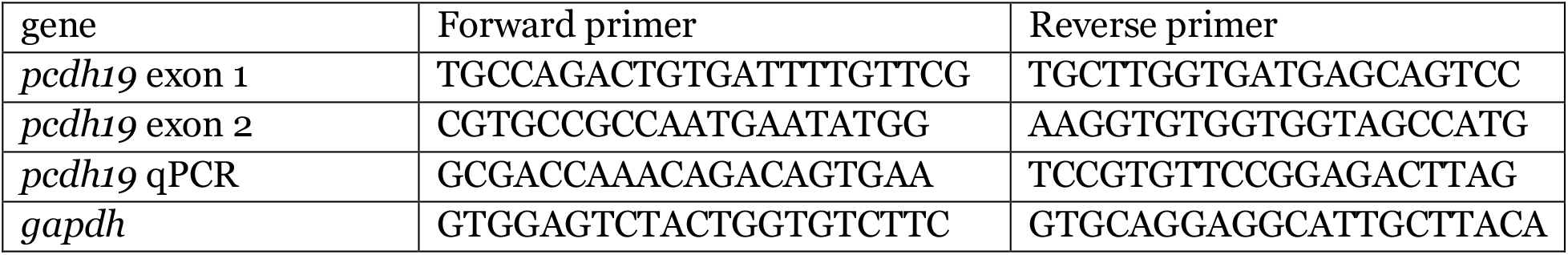
Primers used in PCR and qPCR

### Assessment for spontaneous and induced seizures

At 6dpf larvae with *pcdh19* mutations were transferred into wells (one each) of a 96-well plate, placed into the DanioVision video-tracking system (Noldus, Wageningen, Netherlands), and tracked with the Ethovision software after a 20-minute acclimation period. For 60min, baseline swim patterns were recorded and evaluated for seizure-like episodes.

#### PTZ protocol

Larvae were subjected to a 30min exposure to 2 different concentrations of pentylenetetrazole (PTZ) (0.1mM and 2.5mM).

#### Strobe light protocol

Larvae were subjected to 10min of darkness, three repetitions of strobe light bursts with different frequencies (1Hz for, 2Hz and 5Hz separated by 30sec of darkness) with 12min of light exposure as the final stage of the trial.

#### Hyperthermia protocol

Individual 6dpf larvae were transferred to a 96-well plate submerged in room temperature (22°C) water. Baseline swim patterns were recorded for 20min. Then, the water temperature was slowly elevated over the course of 20min from room temperature to 37°C, and swim patterns were recorded. Afterwards, swim behavior was assessed for seizure-like swim patterns and compared to parameters during the baseline period. As a control for the KO fish lines we used WT animals, and for the mosaic fish we used scrambled gRNA injected animals.

For the analysis, parameters including total distance moved and maximal velocity for each fish were measured in 5-second time bins. Based on these parameters, an automated seizure detection paradigm was developed and used to identify the presence of seizure-like activity in each animal per time bin (Lin et al., 2018).

### Behavioral tests

T-maze cognition test. We designed an asymmetrical T-maze that has one bigger compartment on the right arm versus the left arm. The bigger compartment represents the preference zone (PZ) which is more favorable since its bigger and enriched, which fish are expected to seek out and stay in (Darland and Dowling, 2001). Adult fish (4-5 months old) were released to the maze and are allowed to explore for 5min once a day for 5 consecutive days. The latency from entering the maze to reaching the preference zone as well as the cumulative time the fish spend in the preference zone were analyzed for each fish. Fish that did not move during the whole trial or that did not reach the preference zone were excluded from the analysis. The number of fish that met these exclusion criteria are reported for each genotype.

Bottom dwelling anxiety test. Single, adult, naive fish were placed in a rectangular shaped tank which was virtually divided into upper and lower halves. Their swim behavior was recorded immediately after placement into the new tank for 10min. When exposed to a new environment fish initially seek out the darker, “protective” bottom of a tank to avoid detection by predators and subsequent vertical exploration behavior (Kysil et al., 2017). The higher the anxiety score, the longer the fish stay in the bottom compartment or the more often they will return to the bottom after short episodes of exploring the top compartment. Time spent at the top and bottom-to-top transitions were assessed for each fish.

### Transcriptional analysis

RNA was isolated from zebrafish embryos and larvae. For RNA isolation the tissue from 20 larvae was pooled (to achieve the minimum required RNA concentration), and RNA was then extracted with the QIAGEN RNA isolation kit (QIAGEN, Boston, MA, USA) according to the manufacturers guide. For qPCR, 100ug of RNA was reverse transcribed into cDNA using the SuperScript III (Invitrogen, Waltham, USA) kit according to manufacturer’s instructions. qPCR was performed on a QuantStudio 3 -96-Well 0.1-mL Block and the SYBR green PowerUp kit (Applied Biosystems, Bedford, MA, USA) according to manufacturer’s instructions. Standard curves were obtained for each used primer pair (Table 1), and efficiencies were above 1.8 for each combination.

### LFP recordings

Electrophysiological recordings were obtained from larval zebrafish using previously described methods (Baraban et al., 2013). Larval zebrafish (6 to 7dpf) were paralyzed with alpha-bungarotoxin 2mg/ml (Invitrogen, Waltham, USA), then embedded dorsal side up in 50μl of 1.2% low-melting point agarose (Fisher Scientific, Cambridge, MA, USA) on a recording chamber. The chamber was placed under a Nikon Eclipse FN1 microscope and perfused with zebrafish recording solution containing: 117mM NaCl, 4.7mM KCl, 2.5mM NaHCO_3_, 2.5mM CaCl_2_, 1.2mM MgCl2, 1.2mM NaH_2_PO_4_, 11mM glucose (Sigma-Aldrich, Natick, MA, USA) dissolved in distilled water, brought to a pH of 7.4, and vacuum-filtered through a 0.22μm filter. The specimen was perfused with warm (28.5°C) recording solution at a rate of 1-2ml/min throughout the recording. A borosilicate glass electrode (1-7MΩ resistance) filled with 2M NaCl was placed in the optic tectum using a micromanipulator with visual guidance under the light microscope. Local field potential (LFP) recordings were collected using Clampex software (Molecular Devices, Sunnyvale, USA). All recordings were performed in current-clamp mode, low-pass filtered at 1Hz, high-pass filtered at 1kHz, amplified at a gain of 500x, and sampled at 10kHz (Axopatch 200B amplifier, Digidata 1440A digitizer, Molecular Devices, Sunnyvale, USA). After a baseline recording of 50min, PTZ 40mM (dissolved in recording solution) was added to the recording chamber. The recording continued for 20min after the addition of PTZ. Fish were observed under the microscope frequently throughout the experiment for presence of heartbeat and circulation in the head and absence of movement. Only recordings with a visible heartbeat throughout were included in the analysis.

### Analysis of LFP data

All electrophysiology was analyzed with a custom MATLAB software framework developed by an investigator blind to status of the experiment. The framework first passes the raw data through a filtering stage – based on the Savitzky–Golay filtering algorithm. The software uses the filtered voltages to automatically detect the measurement noise floor by fitting the core of the histogram of the filtered voltages with a Gaussian distribution. This noise floor (0.006 - 0.012mV) allows for a robust detection of all spikes. These spikes are further categorized into small amplitude events (SAE) and large amplitude events (LAE) based on their signal amplitude. This distinction into small and large amplitude event can be directly obtained from a histogram of all detected spike amplitudes in which we observe two distinct features: a narrow peak with low signal amplitude (referred to as the SAE) and a second broad peak with large signal amplitude (referred to as the LAE). The threshold discriminating between small and large amplitude events is 0.08-0.12mV depending on the noise (after completion of the manuscript, the authors became aware of a related classification (Griffin et al., 2021)). A final stage of the software groups the detected large amplitude events into bursts in case the following event occurs within a 1 second time window. This is summarized as the burst frequency, the frequency with which the bursts occur throughout the experiments.

The MATLAB code can be downloaded here: https://github.com/CarstenRobens/EEG_SpikeFinder

### Calcium imaging

*Pcdh19* mutant zebrafish were crossed with Tg(elav3:GCaMP6s); mitfa^nac/nac^ (abbreviated, GCaMP6) zebrafish in the nacre background to achieve GCaMP6-expressing, transparent *pcdh19* mutant lines. Heterozygous *pcdh19* mutants were bred for Ca2+ fluorescence detection at 6dpf.

To this end, individual unrestrained larvae were placed into a well of a 96-well optical plate in 100μl fish water compatible with the Hamamatsu FDSS7000EX fluorescent plate reader. The fish were not evaluated for 20min to allow time for them to acclimate to the new environment. Specimens were illuminated from above via Xenon lamp passed through a 480nm filter. Epifluorescence from below the specimen is passed through a 540nm filter and collected by CCD, allowing all wells to be recorded simultaneously. Data is sampled at approximately 12.7Hz (79msec interval), 16-bit pixel depth and 2×2 binning. Fish were recorded for 30min untreated, 30min treated with 1mM and 30min treated with 10mM PTZ. Afterwards heartbeat is assessed and DNA is extracted for genotype determination. Analysis is performed in MATLAB to extract position, linear and angular velocity, and changes in calcium fluorescence using a moving average ΔF/F_0_ method.

For high-resolution Calcium imaging, living GCaMP6 larvae (5-7dpf) were immobilized with 5mM tubocurare and embedded in 1.2% agarose for imaging using a Zeiss Lightsheet.Z1 microscope equipped with 20X/1.0 NA objective. Calcium fluorescence was stimulated by two co-planar light sheets positioned at 90 degrees from the detection axis using 488nm laser. To assess abnormalities in spontaneous brain activity, two paradigms were used: A) whole brain activity, encompassing ∼13 slices with 22uM slice interval at a rate of 1.8seconds per stack; B) z-section through optic tectum at 10Hz. Samples were immersed in temperature-controlled fish water (28°C) during imaging. Circulation was assessed by transmitted light to confirm vitality of the specimen at the beginning and end of each recording. Analysis was performed in Zen Black software (Zeiss, Jena, Germany).

### Analysis of FDSS data

An algorithm to track changes in calcium activity using a “moving ΔF/F_0_” was devised. The method normalizes the instantaneous average fluorescence for the area of the fish body within the well by the following formula: (average F_fish_(t) - F_0_)/F_0_, where F_0_ = average F_fish_ (averaged over each pixel, for each time sample), and smoothed with a 1000-sample (∼79sec) boxcar moving average. For detecting events, the F/F0 time-series is further smoothed with a 25-sample (∼1.975sec) boxcar moving average. Fish movement is tracked and position, linear velocity, angular velocity estimated. Events are initially detected by identifying peaks that exceeded an empirically determined permissive threshold (>0.05), while the start and end of each event is then identified by the zero-crossing of the smoothed 1st derivative. Subsequently, 47 measurements for each event are obtained related to intensity, distance traveled, linear velocity, angular velocity, total revolutions, and others. Additional measurements for the time-series include total distance, maximum/minimum intensity, and total fish size. Calcium events whose max intensity F/F_0_ exceeded 0.1 were retained as significant.

The code to analyze FDSS calcium imaging data can be downloaded here: https://drive.google.com/file/d/1Gw97gxSiwlE4bFYv7rohNAhvAYUiUixk/view?usp=sharing

### Phosphorylated extracellular signal-related kinase (pERK) staining

pERK staining was performed as described elsewhere with minor modifications (Thyme et al., 2019). Briefly, zebrafish larvae were collected and pooled at 5dpf, given 20min to acclimate in a quiet room and then were quickly fixed in 4% Paraformaldehyde (PFA) + 0.25% Triton X overnight. Pigmentation was removed by bleaching for 10min in sterile fish water containing 3% H2O2, 1% KOH. Samples were washed three times with PBST (PBS with 0.25% Triton X). Antigen retrieval was performed first by incubation of samples in 150mM Tris HCl pH 9.0 for 5min followed by 70°C for 15min. Afterwards, they were washed twice with PBST, for 5min each. The larvae were then permeabilized by incubation in 0.05% Trypsin-EDTA on ice for 45min and subsequently washed three times in PBST. Samples were incubated in blocking solution containing 2% Normal Goat Serum, 1% Bovine Serum Albumin, and 1% DMSO in PBST for 1h. Larvae were incubated with pERK and tERK antibodies in 1% BSA and 1% DMSO in PBST for 48h at 4°C on a rocker. Primary antibodies: mouse anti-total ERK (p44/p42 MAPK (Erk 1/2) (L34F12) mouse mAb Cell Signaling #4696) 1:100, rabbit anti-phospho-ERK (Phospho-p44/42 MAPK (Erk1/2) (Thr202/Tyr204) rabbit mAb Cell Signaling #4370) 1:400. After incubation samples were washed with PBST three times for 15min followed by incubation with the secondary antibody solution (donkey anti-rabbit Alexa Fluor 488 (Invitrogen A21206) 1:200, goat anti-mouse Alexa Fluor 647 (Invitrogen A32728) 1:200) overnight on a rocker at 4°C. Samples were washed three times, 15min each with PBST and stored in PBST at 4°C until imaging. Genotypes were determined after imaging. Analysis of pERK-tERK staining ratio was performed as previously described (Thyme et al., 2019).

### Confocal Imaging

Transgenic Tg(dlx6a-1.4kbdlx5a/dlx6a:GFP::vglut2:DsRed) animals (kindly donated by Ellen Hoffman, Yale School of Medicine, New Haven (Hoffman et al., 2016)) were crossed into the transparent mitfa background and then with our *pcdh19* mutants, resulting in loss of pigmentation and expression of GFP in inhibitory dlx6/5-positve neurons and dsRed in excitatory vGlut2-postive neurons. Adults of this line were allowed to breed for 1h and then all eggs were collected and held at low densities to avoid crowding related maturation delays. Despite the low density, we observed a gradient of maturity in very young larvae, therefore 2dpf larvae used for imaging and analysis were benchmarked based on general developmental milestones (Kimmel et al., 1995) to ensure the use of developmentally matched larvae for experiments. Only hatched larvae that reached at least the long-pec phase with obvious markers of whole-body maturity were included for subsequent brain analysis. Larvae were 4%PFA fixed at 2dpf or 5dpf over night for confocal imaging. Fixed larvae were mounted dorsally in 1.2% agarose and GFP and dsRed fluorescence was imaged immediately after on a Zeiss LSM700 with 15µm step size and a 10x objective lens. For imaging of *pcdh19* mosaic mutants, wildtype transgenic zebrafish were bred and *pcdh19* sgRNAs or scrambled sgRNA was injected into the 1-cell stage.

### TUNEL assay

Zebrafish larvae were fixed overnight in 4% PFA + 0.25% Triton X and washed in PBST. Pigmentation was removed with 10min bleaching in 3% H2O2, 1% KOH in sterile fish water. Samples were incubated in 150mM Tris HCl pH 9.0 for 5min at room temperature, then at 70°C for 15min. Permeabilization was performed in 0.05% Trypsin-EDTA on ice for 45min. Afterwards samples were incubated for 2 h in the In Situ Cell Death Detection Kit (Roche, Basel, Switzerland) enzyme labeling solution at 37°C. Prior to imaging, samples were counterstained in 300nM DAPI for 30min.

### Image Analysis

For cell counting in transgenic *pcdh19* mutants, z-stacks of confocal images were analyzed with genotypes blinded to the user, using ImageJ software. After background subtraction and noise smoothing, a threshold was determined that includes most cells of the tectum and minimizes background fluorescence. Very bright cells were automatically separated in ImageJ using the watershed function. The average number of all cells counted in the tectum of each zebrafish line was reported. Since background noise was substantial in the dsRed-expressing cell populations, we excluded these from the cell counting analysis.

### Western Blot

For each sample 10-20 zebrafish heads from the same previously genotyped genetic background were pooled in RIPA buffer for assessment of Pcdh19 protein levels. Using a tissue homogenizer, larval heads were homogenized for 35-45s on ice, until the samples was uniformly cloudy. Protein was pelleted by centrifugation at 14000rpm for 15min at 4C and resuspended in 20µl buffer for subsequent BCA assay to estimate protein concentration. An equal protein amount was used and mixed with 5μl of 4x Blot sample buffer and DI water and heated for 5min at 95°C. Samples and protein marker were loaded on an SDS-polyacrylamide gel and then blotted on to a PVDF membrane that was pretreated with 100% methanol for 5min and then transferred to 1x transfer buffer. After successful blotting, membranes were blocked in blocking buffer for 1h on a shaker. Primary antibodies were applied in blocking buffer at 4°C over-night (alpha-tubulin, Abcam ab233661, 1:1000 dilution and Pcdh19, Abcam ab191198, in 1:200 dilution). After 3 wash steps in PBS-T (0.1% triton), secondary antibodies IR-Odyssey were applied in blocking buffer in a 1:10000 dilution for 1h at room temperature. Washed membranes were imaged with the LiCor Odyssey system.

### Statistical Analysis

For all experiments *pcdh19* KO lines were compared to WT and *pcdh19* mosaics were compared to scrambled injected (and for LFP recordings in addition to scrambled injected mosaics were also compared to WT and injection controls). All statistical analyses were done using Graphpad Prism 8. Values represent the mean ±SEM. All statistical tests used can be found in the corresponding figure legends. A p-value of <0.05 was determined as statistically significant. N numbers can be found in the corresponding figure legends.

## Acknowledgments

Dr. Barbara Robens is supported by the Deutsche Forschungsgemeinschaft (DFG) Germany research fellowship (#424602704). Dr. McGraw is supported by the CURE Taking Flight Award and NIH/NINDS K08NS118107. Dr. Carsten Robens is supported by the Deutsche Forschungsgemeinschaft (DFG) Germany research fellowship (#421987027). Drs. Rotenberg and Poduri were supported by NIH/NINDS R01NS100766 and the Boston Children’s Hospital Translational Research Program. Dr. Thyme is supported by NIH/NIMH R00MH110603 and a Klingenstein-Simons Foundation Fellowship Award. We thank Dr. Hyun Yong Koh for his support with the transcriptomic data analysis. We are grateful to Seniha Ipekci, Gessica Truglio, Brandon Jones, and other past laboratory members who were involved in the early stages of this project and to the Boston Children’s Hospital Aquatic Facility leadership and staff.

## Author contributions

Conceptualization A.P., B.K.R., experiments performed by B.K.R., X.Y., C.M.M., L.T., S.T., analysis performed by B.K.R., X.Y., C.M.M., C.R., S.T., software written by C.M.M., C.R., manuscript written by B.K.R., A.P.; resources provided by A.P., A.R.; supervision by A.P., B.K.R.

## Supplemental information

**Supplemental Figure 1:**
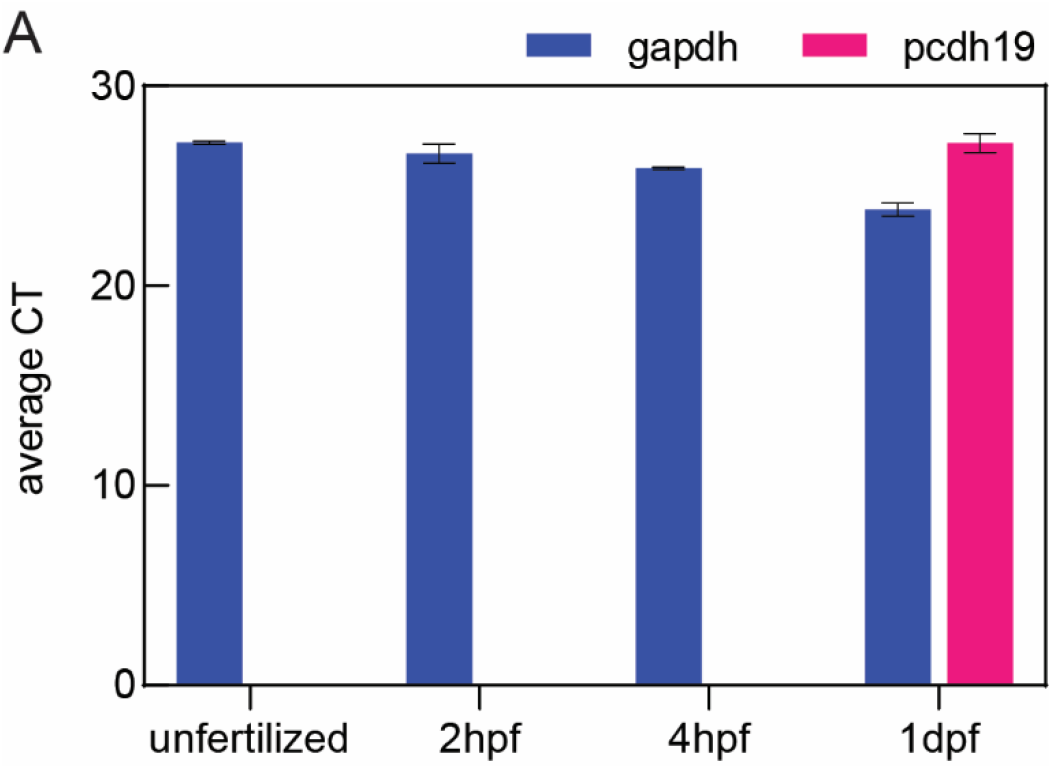
(A) No maternal *pcdh19* mRNA detectable in unfertilized eggs or young embryos. qPCR of WT embryos shows undetectable *pcdh19* mRNA expression at unfertilized, 2hpf, and 4hpf, despite detectable *gapdh* expression. Both *pcdh19* and *gapdh* expression are present at 1dpf. qPCR protocol 40 cycles. N=3 samples of 20 pooled eggs or embryos for each timepoint.

**Supplemental Figure 2:**
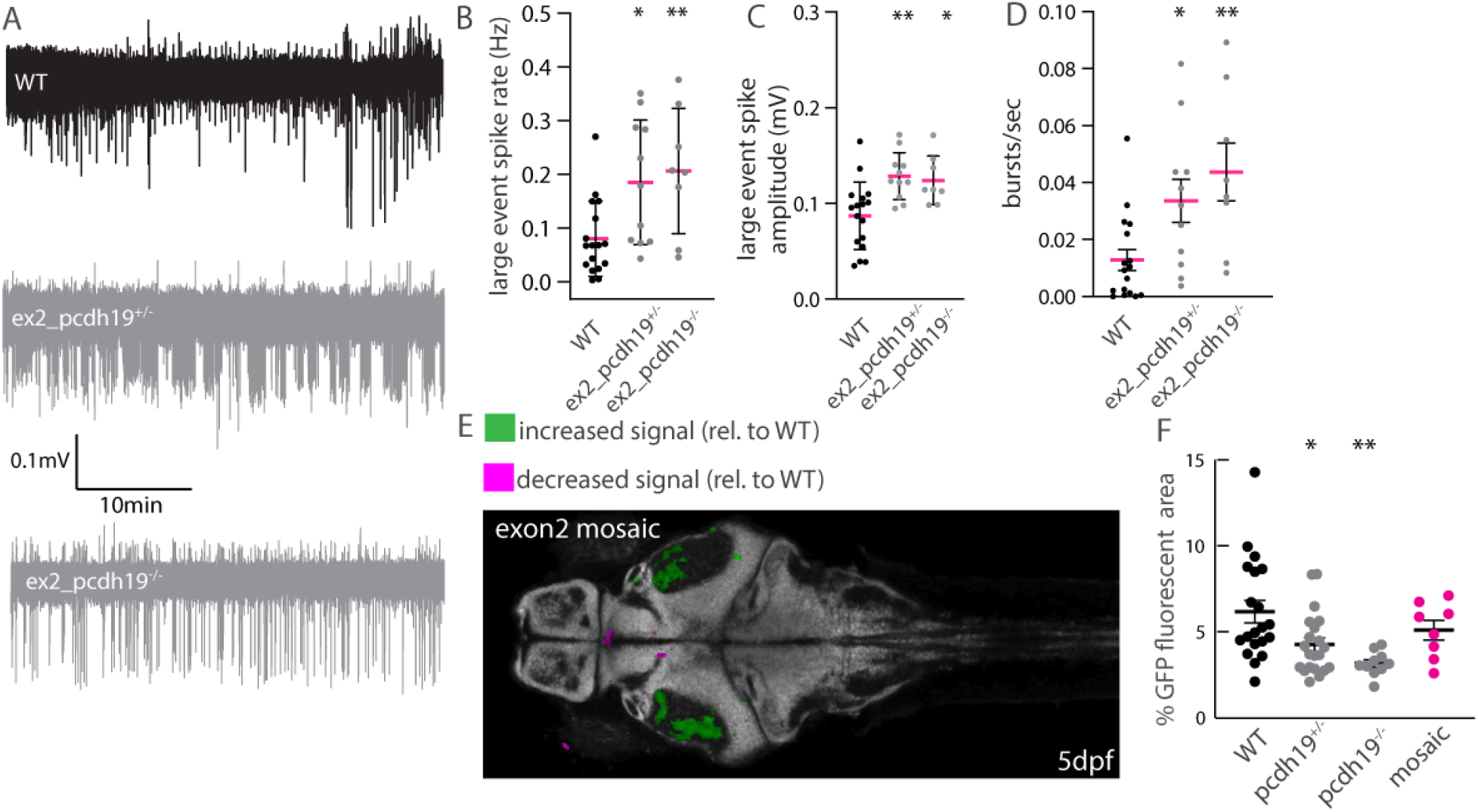
(A) Representative traces of LFP recordings for WT, exon2-pcdh19+/-, and exon2-pcdh19-/-exon2 larvae showing beta events consisting of bursts of spikes present in *pcdh19* mutants and to a lesser extend in controls. (B) Quantification shows a significant increase in beta event rate, (C) in the average beta spike amplitude, and (D) in the average beta burst rate in both exon2 *pcdh19* mutants compared to the WT. n=16 WT, n=11 ex2_*pcdh19*+/-, n=8 ex2_*pcdh19*. One-way ANOVA with Dunnett’s multiple comparisons test, *p<0.05, **p<0.01. (E) Confocal image shows pERK signal intensity is increased within the neuropil of the optic tectum relative to the control. n=23 exon2 mosaic, n=28 scrambled injected larvae. (F) Quantification of the percentage of GFP-positive pixels shows a significant decrease in pcdh19+/- and pcdh19-/-larvae compared to WT and mosaic larvae. n=20 WT, n=21 pcdh19+/-, n=10 pcdh19-/-, n=8 mosaic. One-way ANOVA with Dunnett’s multiple comparisons test, *p<0.05, **p<0.01.

**Supplemental Figure 3:**
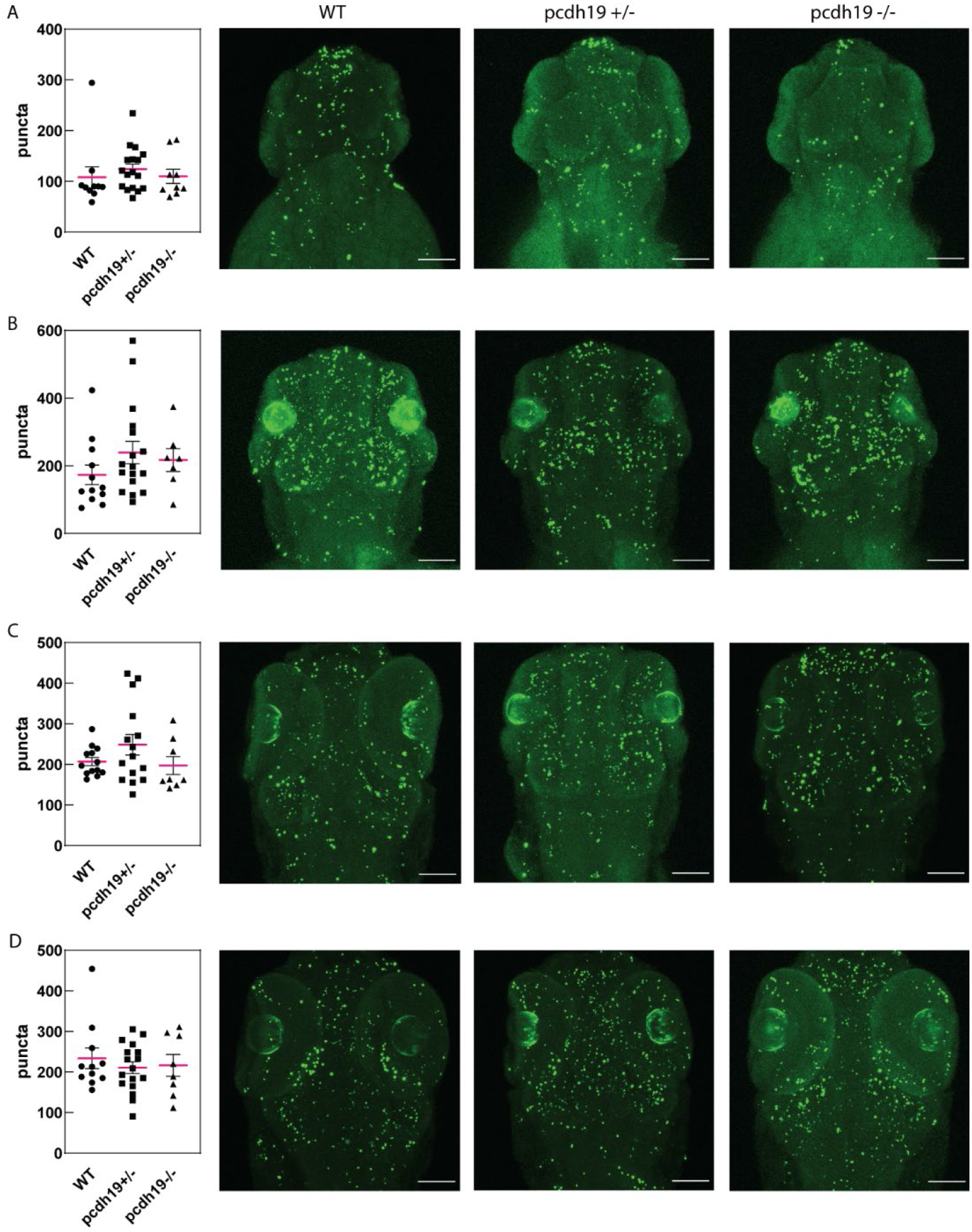
Quantification of TUNEL positive punctae shows no difference in WT compared to pcdh19 KO mutants at (A) 2dpf, (B) 3dpf, (C) 4dpf, and (D) 5dpf. Representative confocal maximum intensity projections of a Z-stack are shown on the right, next to the quantification for each developmental time point and mutant. Scale bar 100µm. 2dpf: 2dpf n=10 WT, n=17 *pcdh19*^+/-^, and n=9 *pcdh19*^-/-^ larvae., 3dpf n=12 WT, n=17 *pcdh19*^+/-^, and n=7 *pcdh19*^-/-^ larvae, 4dpf: n=13 WT, n=15 *pcdh19*^+/-^, and n=8 *pcdh19*^-/-^ larvae, 5dpf n=11 WT, n=17 *pcdh19*^+/-^, and n=8 *pcdh19*^-/-^ larvae. One-way ANOVA with Tukey’s multiple comparisons test.

**Supplemental video 1**: Light sheet calcium imaging video of a 7dpf pcdh19^+/-^ larvae with spontaneous calcium activity

**Supplemental video 2 & 3**: FDSS low-resolution calcium imaging video clip of a heterozygous pcdh19 mutant having a seizure and a calcium event.

## References

Afrikanova, T., Serruys, A.S., Buenafe, O.E., Clinckers, R., Smolders, I., de Witte, P.A., Crawford, A.D., and Esguerra, C.V. (2013). Validation of the zebrafish pentylenetetrazol seizure model: locomotor versus electrographic responses to antiepileptic drugs. PLoS One 8, e54166. 10.1371/journal.pone.0054166

Ashburner, M., Ball, C.A., Blake, J.A., Botstein, D., Butler, H., Cherry, J.M., Davis, A.P., Dolinski, K., Dwight, S.S., Eppig, J.T., Harris, M.A., Hill, D.P., Issel-Tarver, L., Kasarskis, A., Lewis, S., Matese, J.C., Richardson, J.E., Ringwald, M., Rubin, G.M., and Sherlock, G. (2000). Gene ontology: tool for the unification of biology. The Gene Ontology Consortium. Nat Genet 25, 25–29. 10.1038/75556

Baraban, S.C., Dinday, M.T., and Hortopan, G.A. (2013). Drug screening in Scn1a zebrafish mutant identifies clemizole as a potential Dravet syndrome treatment. Nat Commun 4, 2410. 10.1038/ncomms3410

Bassani, S., Cwetsch, A.W., Gerosa, L., Serratto, G.M., Folci, A., Hall, I.F., Mazzanti, M., Cancedda, L., and Passafaro, M. (2018). The female epilepsy protein PCDH19 is a new GABAAR-binding partner that regulates GABAergic transmission as well as migration and morphological maturation of hippocampal neurons. Hum Mol Genet 27, 1027–1038. 10.1093/hmg/ddy019

Blevins, C.J., Emond, M.R., Biswas, S., and Jontes, J.D. (2011). Differential expression, alternative splicing, and adhesive properties of the zebrafish delta1-protocadherins. Neuroscience 199, 523–534. 10.1016/j.neuroscience.2011.09.061

Cooper, S.R., Emond, M.R., Duy, P.Q., Liebau, B.G., Wolman, M.A., and Jontes, J.D. (2015). Protocadherins control the modular assembly of neuronal columns in the zebrafish optic tectum. J Cell Biol 211, 807–814. 10.1083/jcb.201507108

Darland, T., and Dowling, J.E. (2001). Behavioral screening for cocaine sensitivity in mutagenized zebrafish. Proc Natl Acad Sci U S A 98, 11691–11696. 10.1073/pnas.191380698

Dentici, M.L., Alesi, V., Quinodoz, M., Robens, B., Guerin, A., Lebon, S., Poduri, A., Travaglini, L., Graziola, F., Afenjar, A., Keren, B., Licursi, V., Capuano, A., Dallapiccola, B., Superti-Furga, A., and Novelli, A. (2021). Biallelic variants in ZNF526 cause a severe neurodevelopmental disorder with microcephaly, bilateral cataract, epilepsy and simplified gyration. J Med Genet. 10.1136/jmedgenet-2020-107430

Depienne, C., Bouteiller, D., Keren, B., Cheuret, E., Poirier, K., Trouillard, O., Benyahia, B., Quelin, C., Carpentier, W., Julia, S., Afenjar, A., Gautier, A., Rivier, F., Meyer, S., Berquin, P., Helias, M., Py, I., Rivera, S., Bahi-Buisson, N., Gourfinkel-An, I., Cazeneuve, C., Ruberg, M., Brice, A., Nabbout, R., and Leguern, E. (2009). Sporadic infantile epileptic encephalopathy caused by mutations in PCDH19 resembles Dravet syndrome but mainly affects females. PLoS Genet 5, e1000381. 10.1371/journal.pgen.1000381

Depienne, C., and LeGuern, E. (2012). PCDH19-related infantile epileptic encephalopathy: an unusual X-linked inheritance disorder. Human mutation 33, 627–634. 10.1002/humu.22029

Dibbens, L.M., Tarpey, P.S., Hynes, K., Bayly, M.A., Scheffer, I.E., Smith, R., Bomar, J., Sutton, E., Vandeleur, L., Shoubridge, C., Edkins, S., Turner, S.J., Stevens, C., O’Meara, S., Tofts, C., Barthorpe, S., Buck, G., Cole, J., Halliday, K., Jones, D., Lee, R., Madison, M., Mironenko, T., Varian, J., West, S., et al. (2008). X-linked protocadherin 19 mutations cause female-limited epilepsy and cognitive impairment. Nat Genet 40, 776–781. 10.1038/ng.149

Dimova, P.S., Kirov, A., Todorova, A., Todorov, T., and Mitev, V. (2012). A novel PCDH19 mutation inherited from an unaffected mother. Pediatric neurology 46, 397–400. 10.1016/j.pediatrneurol.2012.03.004

Duszyc, K., Terczynska, I., and Hoffman-Zacharska, D. (2015). Epilepsy and mental retardation restricted to females: X-linked epileptic infantile encephalopathy of unusual inheritance. J Appl Genet 56, 49–56. 10.1007/s13353-014-0243-8

Eimon, P.M., Ghannad-Rezaie, M., De Rienzo, G., Allalou, A., Wu, Y., Gao, M., Roy, A., Skolnick, J., and Yanik, M.F. (2018). Brain activity patterns in high-throughput electrophysiology screen predict both drug efficacies and side effects. Nat Commun 9, 219. 10.1038/s41467-017-02404-4

Epi4K Consortium, and Epilepsy Phenome/Genome Project (2017). Ultra-rare genetic variation in common epilepsies: a case-control sequencing study. Lancet Neurol 16, 135–143. 10.1016/S1474-4422(16)30359-3

Griffin, A., Carpenter, C., Liu, J., Paterno, R., Grone, B., Hamling, K., Moog, M., Dinday, M.T., Figueroa, F., Anvar, M., Ononuju, C., Qu, T., and Baraban, S.C. (2021). Phenotypic analysis of catastrophic childhood epilepsy genes. Commun Biol 4, 680. 10.1038/s42003-021-02221-y

Griffin, A., Hamling, K.R., Knupp, K., Hong, S., Lee, L.P., and Baraban, S.C. (2017). Clemizole and modulators of serotonin signalling suppress seizures in Dravet syndrome. Brain 140, 669–683. 10.1093/brain/aww342

Grone, B.P., Marchese, M., Hamling, K.R., Kumar, M.G., Krasniak, C.S., Sicca, F., Santorelli, F.M., Patel, M., and Baraban, S.C. (2016). Epilepsy, Behavioral Abnormalities, and Physiological Comorbidities in Syntaxin-Binding Protein 1 (STXBP1) Mutant Zebrafish. PLoS One 11, e0151148. 10.1371/journal.pone.0151148

Hayashi, S., Inoue, Y., Hattori, S., Kaneko, M., Shioi, G., Miyakawa, T., and Takeichi, M. (2017). Loss of X-linked Protocadherin-19 differentially affects the behavior of heterozygous female and hemizygous male mice. Sci Rep 7, 5801. 10.1038/s41598-017-06374-x

Hoffman, E.J., Turner, K.J., Fernandez, J.M., Cifuentes, D., Ghosh, M., Ijaz, S., Jain, R.A., Kubo, F., Bill, B.R., Baier, H., Granato, M., Barresi, M.J., Wilson, S.W., Rihel, J., State, M.W., and Giraldez, A.J. (2016). Estrogens Suppress a Behavioral Phenotype in Zebrafish Mutants of the Autism Risk Gene, CNTNAP2. Neuron 89, 725–733. 10.1016/j.neuron.2015.12.039

Homan, C.C., Pederson, S., To, T.H., Tan, C., Piltz, S., Corbett, M.A., Wolvetang, E., Thomas, P.Q., Jolly, L.A., and Gecz, J. (2018). PCDH19 regulation of neural progenitor cell differentiation suggests asynchrony of neurogenesis as a mechanism contributing to PCDH19 Girls Clustering Epilepsy. Neurobiol Dis 116, 106–119. 10.1016/j.nbd.2018.05.004

Hortopan, G.A., Dinday, M.T., and Baraban, S.C. (2010). Spontaneous seizures and altered gene expression in GABA signaling pathways in a mind bomb mutant zebrafish. J Neurosci 30, 13718–13728. 10.1523/JNEUROSCI.1887-10.2010

Hoshina, N., Johnson-Venkatesh, E.M., Hoshina, M., and Umemori, H. (2021). Female-specific synaptic dysfunction and cognitive impairment in a mouse model of PCDH19 disorder. Science 372. 10.1126/science.aaz3893

Hunt, R.F., Hortopan, G.A., Gillespie, A., and Baraban, S.C. (2012). A novel zebrafish model of hyperthermia-induced seizures reveals a role for TRPV4 channels and NMDA-type glutamate receptors. Exp Neurol 237, 199–206. 10.1016/j.expneurol.2012.06.013

Kimmel, C.B., Ballard, W.W., Kimmel, S.R., Ullmann, B., and Schilling, T.F. (1995). Stages of embryonic development of the zebrafish. Dev Dyn 203, 253–310. 10.1002/aja.1002030302

Kolc, K.L., Sadleir, L.G., Scheffer, I.E., Ivancevic, A., Roberts, R., Pham, D.H., and Gecz, J. (2018). A systematic review and meta-analysis of 271 PCDH19-variant individuals identifies psychiatric comorbidities, and association of seizure onset and disease severity. Mol Psychiatry. 10.1038/s41380-018-0066-9

Kysil, E.V., Meshalkina, D.A., Frick, E.E., Echevarria, D.J., Rosemberg, D.B., Maximino, C., Lima, M.G., Abreu, M.S., Giacomini, A.C., Barcellos, L.J.G., Song, C., and Kalueff, A.V. (2017). Comparative Analyses of Zebrafish Anxiety-Like Behavior Using Conflict-Based Novelty Tests. Zebrafish 14, 197–208. 10.1089/zeb.2016.1415

Lin, X., Duan, X., Jacobs, C., Ullmann, J., Chan, C.Y., Chen, S., Cheng, S.H., Zhao, W.N., Poduri, A., Wang, X., Haggarty, S.J., and Shi, P. (2018). High-throughput brain activity mapping and machine learning as a foundation for systems neuropharmacology. Nat Commun 9, 5142. 10.1038/s41467-018-07289-5

Liu, J., and Baraban, S.C. (2019). Network Properties Revealed during Multi-Scale Calcium Imaging of Seizure Activity in Zebrafish. eNeuro 6. 10.1523/ENEURO.0041-19.2019

Meeker, N.D., Hutchinson, S.A., Ho, L., and Trede, N.S. (2007). Method for isolation of PCR-ready genomic DNA from zebrafish tissues. Biotechniques 43, 610, 612, 614. 10.2144/000112619

Mincheva-Tasheva, S., Nieto Guil, A.F., Homan, C.C., Gecz, J., and Thomas, P.Q. (2021). Disrupted Excitatory Synaptic Contacts and Altered Neuronal Network Activity Underpins the Neurological Phenotype in PCDH19-Clustering Epilepsy (PCDH19-CE). Mol Neurobiol 58, 2005–2018. 10.1007/s12035-020-02242-4

Pederick, D.T., Homan, C.C., Jaehne, E.J., Piltz, S.G., Haines, B.P., Baune, B.T., Jolly, L.A., Hughes, J.N., Gecz, J., and Thomas, P.Q. (2016). Pcdh19 Loss-of-Function Increases Neuronal Migration In Vitro but is Dispensable for Brain Development in Mice. Sci Rep 6, 26765. 10.1038/srep26765

Pederick, D.T., Richards, K.L., Piltz, S.G., Kumar, R., Mincheva-Tasheva, S., Mandelstam, S.A., Dale, R.C., Scheffer, I.E., Gecz, J., Petrou, S., Hughes, J.N., and Thomas, P.Q. (2018). Abnormal Cell Sorting Underlies the Unique X-Linked Inheritance of PCDH19 Epilepsy. Neuron 97, 59–66 e55. 10.1016/j.neuron.2017.12.005

Rakotomamonjy, J., Sabetfakhri, N.P., McDermott, S.L., and Guemez-Gamboa, A. (2020). Characterization of seizure susceptibility in Pcdh19 mice. Epilepsia 61, 2313–2320. 10.1111/epi.16675

Samanta, D. (2020). PCDH19-Related Epilepsy Syndrome: A Comprehensive Clinical Review. Pediatr Neurol 105, 3–9. 10.1016/j.pediatrneurol.2019.10.009

Schnorr, S.J., Steenbergen, P.J., Richardson, M.K., and Champagne, D.L. (2012). Measuring thigmotaxis in larval zebrafish. Behav Brain Res 228, 367–374. 10.1016/j.bbr.2011.12.016

Smith, L., Singhal, N., El Achkar, C.M., Truglio, G., Rosen Sheidley, B., Sullivan, J., and Poduri, A. (2018). PCDH19-related epilepsy is associated with a broad neurodevelopmental spectrum. Epilepsia. 10.1111/epi.14003

Tan, C., Shard, C., Ranieri, E., Hynes, K., Pham, D.H., Leach, D., Buchanan, G., Corbett, M., Shoubridge, C., Kumar, R., Douglas, E., Nguyen, L.S., McMahon, J., Sadleir, L., Specchio, N., Marini, C., Guerrini, R., Moller, R.S., Depienne, C., Haan, E., Thomas, P.Q., Berkovic, S.F., Scheffer, I.E., and Gecz, J. (2015). Mutations of protocadherin 19 in female epilepsy (PCDH19-FE) lead to allopregnanolone deficiency. Hum Mol Genet 24, 5250–5259. 10.1093/hmg/ddv245

Thyme, S.B., Pieper, L.M., Li, E.H., Pandey, S., Wang, Y., Morris, N.S., Sha, C., Choi, J.W., Herrera, K.J., Soucy, E.R., Zimmerman, S., Randlett, O., Greenwood, J., McCarroll, S.A., and Schier, A.F. (2019). Phenotypic Landscape of Schizophrenia-Associated Genes Defines Candidates and Their Shared Functions. Cell 177, 478–491 e420. 10.1016/j.cell.2019.01.048

Turrini, L., Fornetto, C., Marchetto, G., Mullenbroich, M.C., Tiso, N., Vettori, A., Resta, F., Masi, A., Mannaioni, G., Pavone, F.S., and Vanzi, F. (2017). Optical mapping of neuronal activity during seizures in zebrafish. Sci Rep 7, 3025. 10.1038/s41598-017-03087-z

Wang, W., and Frankel, W.N. (2021). Overlaps, gaps, and complexities of mouse models of Developmental and Epileptic Encephalopathy. Neurobiol Dis 148, 105220. 10.1016/j.nbd.2020.105220

Wu, Q. (2005). Comparative genomics and diversifying selection of the clustered vertebrate protocadherin genes. Genetics 169, 2179–2188. 10.1534/genetics.104.037606

