## Supplementary figures and images for "Mosaic and non-mosaic protocadherin 19 mutation leads to neuronal hyperexcitability in zebrafish"

### Supplementary video 2

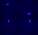

### Supplementary video 3

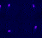
